# Summing the parts: Improving population estimates using a state-space multispecies production model

**DOI:** 10.1101/2025.02.20.639295

**Authors:** Paul M. Regular, Mariano Koen-Alonso, M. Joanne Morgan, Pierre Pepin, Rick M. Rideout

**Affiliations:** Fisheries and Oceans Canada, Northwest Atlantic Fisheries Centre, 80 East White Hills, St. John’s, Newfoundland and Labrador, A1C 5X1, Canada

## Abstract

Carrying capacity is a fundamental concept in ecology that has inspired the development and application of a broad range of population models. In the context of fisheries science, production models have been employed globally to calculate carrying capacity and guide the sustainable use of fish populations. Production models have, however, been criticized for failing to account for species interactions and environmental effects. We aim to fill some of these gaps by introducing a novel state-space multispecies production model. We apply our extended model to commercially important demersal fish species off the east coast of Canada to assess its ability to reveal species interactions and the relative impacts of fishing and environmental effects. Our results indicate that accounting for species interactions increases the accuracy of biomass estimates for species within a community. The model also revealed strongly correlated process deviations, unrelated to fishing or density-dependent effects, which unexpectidly indicates that widespread collapses were primarily driven by a common environmental driver rather than fishing. Such inferences indicate that this may be a promising avenue for producing more holistic and accurate assessments for multiple species with relatively minimal data requirements (time-series of landings and fisheries-independent indices). Finally, this approach may serve as a stepping stone towards an ecosystem-based approach to fisheries management.

## Introduction

The concept of carrying capacity has long been foundational in applied population ecology, being widely used in the management of renewable resources (Chapman & Byron, 2018; Hilborn et al., 1995). The understanding that populations produce more offspring than an environment can sustain led to the notion that the ‘surplus’ can be harvested sustainably (Pauly & Froese, 2021). These ideas are exemplified by the use of maximum sustainable yield (MSY) in the classic single-species surplus production modeling framework (Schaefer, 1954). In one equation, this framework attempts to explain interannual changes in biomass using fisheries landings and estimates of intrinsic growth rate and carrying capacity. Given the theoretical elegance of the approach, it has been both widely adopted and scrutinized. Estimates of carrying capacity, and resultant derivations of MSY, are frequently criticized for being time-invariant, which ignores the ubiquity of environmental variation (Del Monte-Luna et al., 2004). Moreover, traditional surplus production models tend to focus on single-species dynamics and often disregard the complexities arising from species interactions within ecosystems (Gamble & Link, 2009).

Recognizing the limitations of single-species approaches, there have been calls to move towards an ecosystem-based approach to fisheries management (EBFM; Latour et al., 2003). EBFM acknowledges the intricate web of ecological interactions and aims to ensure the sustainability and integrity of marine ecosystems while supporting viable fisheries (Pikitch et al., 2004). To successfully implement EBFM, it is crucial to develop models that account for species interactions and the dynamics of multiple species within the ecosystem. Substantial progress has been made in the development of multispecies models and a spectrum of approaches have been developed, ranging from complex models that attempt to account for all parts of marine ecosystems (e.g., Fulton et al., 2011) to multispecies age-structured assessment models (e.g., Albertsen et al., 2018) to multispecies surplus production models (e.g., Bundy et al., 2012; Gamble & Link, 2009; Mueter & Megrey, 2006). However, the application of these approaches to fisheries management have often been hindered by data limitations (e.g., age data are frequently not available) and knowledge gaps (e.g., incomplete understanding of food-web interactions). There is therefore a need for methods to help bridge the gap between single-species and multispecies assessment in data or information poor systems.

In this paper, we borrow concepts from single-species surplus production modeling (Millar & Meyer, 2000) and multispecies modeling (Albertsen et al., 2018) to construct a model that incorporates the impacts of fishing on single-species populations and accounts for species interactions within the ecosystem. The data requirements of this model are relatively minimal, requiring species-specific landings and survey indices of biomass. As a case study, we apply this model to commercially important demersal fish species off the east coast of Canada, specifically the Grand Banks of Newfoundland. This case study is particularly germane due to the widespread collapse of most stocks in the area during the early 1990s (Lear, 1998), leaving the relative contributions of fishing and environmental impacts uncertain (Pedersen et al., 2017). Using this case study, we aim to reveal species interactions, distinguish the impacts of fishing from environmental effects, and demonstrate the utility of this modeling approach. The subsequent sections of this paper will present the conceptual framework of our model, describe its application, and discuss the implications of our findings for ecosystem-based fisheries management.

## Methods

### Model formulation

Trends in fish populations have frequently been described using state-space production models of the form

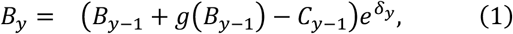

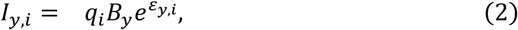

where *B*_*y*_ is biomass at the start of year *y, C*_*y*_ is the catch through year *y, g*(*B*) is production as a function of biomass, *I*_*y,s*_ is an index of relative abundance in year *y* from survey *i, q*_*i*_ is the time-invariant catchability coefficient for survey index *i, δ* is process error, and *ε* is observation error. Statistical challenges aside, the most difficult aspect of this model to parameterize is the production function as it needs to capture changes caused by growth, recruitment, and natural mortality. Schaefer (1954) proposed a solution by applying the logistic equation to describe self-limiting growth,

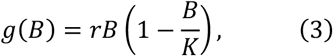

where *r* is the maximum per-capita rate of change and *K* is the carrying capacity. That is, a populations’ intrinsic ability to grow (*rB*) is limited by the size of the current population relative to the maximum biomass the system can support (1 − *B*/*K*). While this formulation offers an elegant description of single-species population dynamics, it assumes that density-dependent effects are solely caused by intraspecific competition and ignores the potential effects of other species inhabiting the same ecological area, competing for the same resources. We present an extension of equation (3) that attempts to account for intra and interspecific competition by assuming that density-dependent effects are incurred when the total biomass of multiple species, represented by *s*, exceeds the capacity of the system, *K*_Σ_,

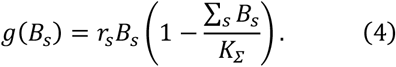

While intrinsic rates of growth may vary across species, this formulation implies that the growth of all species is ultimately limited by the finite amount of energy in a region (i.e., as the total population of all species in the system increases towards *K*_Σ_, year-over-year growth of all species slows). Combining equations (1), (2), and (4), our model becomes

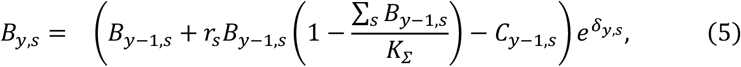

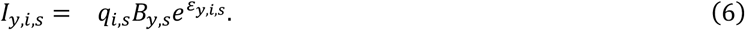

The inclusion of multiple species in the model permits the estimation of covariance. While covarying changes may be apparent in the observations, we assume that most covariance stems from ecosystem processes. We therefore treat observation errors as independent and normally distributed deviations such that *ε*_*y,i,s*_ ∼ *N*(0, τ_*i,s*_), where the standard deviation parameter τ_*i,s*_ represents species and survey specific levels of observation error. A more flexible error structure is used to describe the process errors to account for temporal dependencies driven by as ecological processes. For instance, species interactions may drive positive or negative population responses resulting from direct or indirect associations. Deviations from expected production may also display temporal dependence if the factors contributing to the process errors change gradually over time. Such inertia may cause positive or negative process errors to persist across several years. A first-order autoregressive (AR1) process was therefore applied to account for temporal dependence. Both sources of dependence are modeled using a multivariate AR1 process where

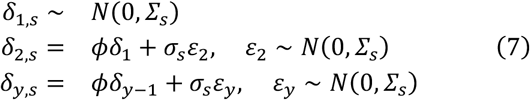

and

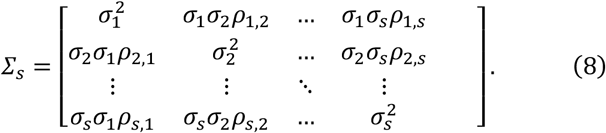

The degree of temporal correlation is controlled by *ϕ*, where values between 0 to 1 represent low to high correlation, and species-to-species correlations are described by *ρ*_*s,s*_, where values between -1 to 1 represent negative to positive correlation. This is a flexible structure that allows for the testing of alternate hypotheses that process errors are independent through time or across species (i.e, *ϕ* = 0 or *ρ*_*s,s*_ = 0). The possibility that process errors are similarly correlated across all species may also be tested by estimating only one *ρ* parameter. Finally, the magnitude of the process error deviations is controlled by the species-specific standard deviation parameters, *σ*_*s*_.

Minor extensions of the formulation also permit the fitting of covariates which may describe an underlying linear effect. Two options were implemented, one that affects the process errors by substituting *e*^*δy,s*^ in equation (5) with 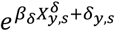, and another that affects the carrying capacity by substituting *K*_Σ_ in equation (5) with 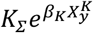, where *β* parameters capture linear effects of covariates included in design matrices, *X*. The idea is that some factors may affect positive or negative changes in the populations while others may affect change in the total carrying capacity of the system. A covariate option for intrinsic rates of increase was not implemented since one goal of this model is to obtain estimates of this vital rate, which is not expected to change rapidly as it is shaped by natural selection (Hutchings, 2021). The formulation was also modified to fit the single-species Schaefer production function by dropping the summation of biomass in equation (5) and estimating species-specific carrying capacities, *K*_*s*_, rather than a system level carrying capacity, *K*_Σ_ (i.e., apply equation (3) indexed by species).

### Statistical framework

This model was implemented using template model builder (TMB; Kristensen et al., 2016), which is a R (R Core Team, 2021) package that enables the fitting of complex nonlinear random effects such as the latent *B* variable in state-space production models (equation (1)). Such variables are not directly measured but are inferred indirectly via observed values. Data fitting is accomplished using a combination of Laplace approximation and automatic differentiation to evaluate the joint likelihood (Kristensen et al., 2016). Like the production model described by Pedersen et al. (2017), both frequentist and Bayesian inference of model parameters are possible. In development, we found that estimation of parameters was generally more successful when vaguely informative priors were specified, and in some cases, not estimatable when unconstrained.

### Case study

#### Data

The multispecies production model described above requires two basic inputs for each species: a time-series of catch (*C*_*y,s*_ in equation (5)), and an index of population size (*I*_*y,s*_ in equation (6)). The Northwest Atlantic Fisheries Organization (NAFO) and Fisheries and Oceans Canada (DFO) have been collecting and curating such information for multiple fish populations along the shelves of Newfoundland and Labrador (NL) since the 1970s. The communities inhabiting these shelves can be divided into several regions with distinct productivity [i.e. ecosystem production units; Pepin et al. (2014)]. For our case study, we tallied catch data and calculated survey indices of multiple demersal fish populations from three regions (Figure 1): 1) the Northeast NL Shelf (NAFO divisions 2J3K), 2) the Grand Bank (NAFO divisions 3LNO), and 3) Southern NL (NAFO sub-division 3Ps).

**Fig 1:**
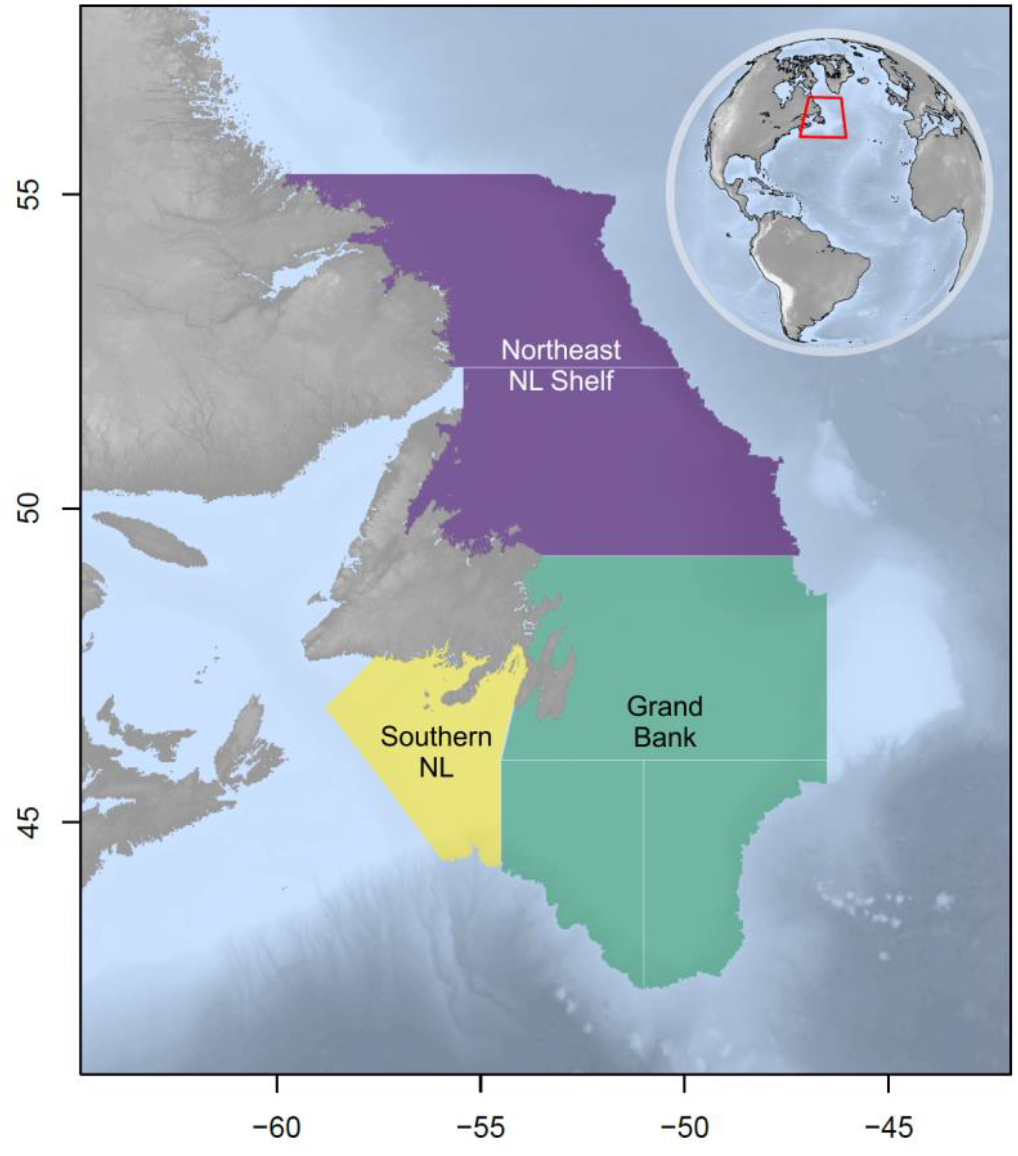
Map of Newfoundland and Labrador (NL) case study area showing the Northeast NL Shelf (purple), Grand Bank (green), and Southern NL (yellow) ecosystem production units.

Catch data were extracted from NAFO’s STATLANT 21A database (https://www.nafo.int/Data/STATLANT-21A, accessed 2022-01-21) and aggregated by region, species, and year. Survey indices were derived from the standardized, stratified random bottom-trawl surveys conducted each spring and fall by DFO; this is perhaps the largest fisheries-independent survey conducted in the world, which aims to cover more than 500,000 km^2^ annually (roughly the size of Sweden or the Yukon, Canada) to depth up to 1500 m. Since the inception of this program in 1971, survey platforms and protocols have undergone a series of changes that affect the continuity of the data collected in each region and season. A Yankee then Engel otter trawl, with nets designed to catch large demersal fish, were used between 1971 to 1994. Starting in the fall of 1995 survey gear was changed to a Campelen shrimp trawl with a small mesh codend, which allowed a broader range of species and size groups to be captured (Chadwick et al., 2007). Within each era of the survey (Yankee, Engel, or Campelen) and for each season and region, samples used in this study were limited to strata that were covered most years (> 80%) and to species found across more than 10% of these core strata. This often resulted in the exclusion of strata >750 m as these areas have been inconsistently covered by the survey. Stratified analyses (Smith & Somerton, 1981) were then conducted on the remaining species to obtain indices of total biomass. To minimize bias introduced by inconsistent survey coverage, indices from years where more than 20% of the biomass was likely missed, inferred from time averaged percent occupancy within strata, were excluded from our analysis [*sensu* NAFO guidelines; page 10, NAFO (2019)]. Finally, species were ranked by cumulative commercial catch and limited to the seven most commonly caught species, or species group when catch was not consistently distinguished by species, within each region. On the Northeast NL Shelf, the included species were Redfish spp. (*Sebastes fasciatus* and *Sebastes mentella* combined), Wolffish spp. (*Anarhichas lupus* and *Anarhichas minor* combined), Witch Flounder (*Glyptocephalus cynoglossus*), American Plaice (*Hippoglossoides platessoides*), Greenland Halibut (*Reinhardtius hippoglossoides*), Atlantic Cod (*Gadus morhua*), and Skate spp. (*Amblyraja radiata* and *Malacoraja senta* combined). On the Grand Bank, the included species were Redfish spp., Yellowtail Flounder (*Myzopsetta ferruginea*), American Plaice, Greenland Halibut, Haddock (*Melanogrammus aeglefinus*), and Atlantic Cod. Finally, along Southern NL, the species included were Redfish spp., Witch Flounder, American Plaice, White hake (*Urophycis tenuis*), Haddock, Atlantic Cod, and Skate spp.

#### Priors

For simplicity, all priors were normally distributed and, in most cases, upper and lower inflection points (*μ* ± *σ*; ∼68% of the total area under the curve) of each normal prior were defined using values in log or logit space. Upper and lower values were based on previous research or knowledge to impose fairly generic and vaguely informative priors.

##### Intrinsic growth rate, r_s_, and carrying capacity, K

To capture a broad range of intrinsic growth rates (Thorson, 2020), 0.01 and 1 were chosen as the lower, *r*_*l*_, and upper, *r*_*u*_, values; this translates to a normal prior with a mean of -1.15 and a standard deviation of 1.15 on the log-scale. The log-scale *K*_Σ_ prior was informed by total levels of catch and *r*_*l*_ and *r*_*u*_,

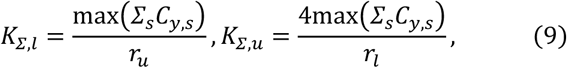

where *K*_Σ,*l*_ and *K*_Σ,*u*_ are the lower and upper values. On the lower end, this prior imposes the assumption that the fishery is unlikely to have caught the equivalent of *K*_Σ_ and, on the upper end, it assumes that the maximum observed catch represents a portion of *K*_Σ_ (*sensu* Froese et al., 2017). The division by the lower and upper values for *r* also accounts for the potential range of productivity. Note that the maximum time-series catches are species specific when estimating species-specific carrying capacities.

##### *Starting biomass, B*_1,*s*_

Like *K*_Σ_, catch was used to constrain the plausible range of the biomass at the beginning of the time series, *B*_1,*s*_, which we will denote as *B*0 to simplify notation. Specifically,

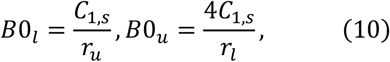

where *B*0_*l*_ and *B*0_*u*_ are the lower and upper values that are log transformed to define the normal prior for log(*B*0). Again, adjusting for the potential range of productivity from *r*_*l*_ to *r*_*u*_, this assumes that the fishery did not catch all of the biomass in the first year and it assumes that the catch represents a portion of the biomass in the first year. A potential flaw with the upper value for this prior is that a lack of market demand may contradict the assumption that landings are coarsely proportional to stock size. However, the upper range chosen was considered reasonable for this case study as there was an active fishery for each species examined here at the start of the time series.

##### Process error variance, 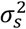, temporal correlation, ϕ, and species-to-species correlation, ρ_s,s_

Considerable process variability may arise from variable recruitment, natural mortality, and/or growth. For instance, species in the Scorpaenidae family, which includes the *Sebastes* genus, have notoriously variable recruitment which frequently results in “spasmodic” stock dynamics (Cadigan et al., 2022; Licandeo et al., 2020). Also, there is evidence that heightened and variable natural mortality contributed to the collapse and slow recovery of Atlantic cod along the Northeast NL Shelf (Cadigan, 2015; Regular et al., 2022). To account for a wide range of possible standard deviations, *σ*_*s*_, a vague prior with 0.01 and 1 as the lower and upper values were chosen. In log-space, this translates to a normal distribution with a mean of -2.3 and sd of 2.3.

There is little knowledge to determine the potential level of temporal or species-to-species dependence present in the focal systems; vague priors were therefore defined for *ϕ* and *ρ*_*s,s*_. For *ϕ*, 0.1 and 0.9 were logit transformed [log(*x*/(1 − *x*))] to define the upper and lower inflection points for a normal prior for the logit of *ϕ*. For *ρ*, the logit transformation was shifted [log((1 *+ x*)/(1 − *x*))] to capture negative and positive lower (−0.9) and upper (0.9) values. When these parameters are estimated, these priors give most credence to moderate temporal correlation (0.5) and no species-to-species correlation (0) but still allow the possibility of high levels of correlation (0.9).

##### Catchability, q_i,s_, and observation error variance,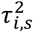

The catchability of the spring and fall surveys conducted by DFO likely changed over time given the shift in gear from a Yankee to Engel to Campelen trawl as well as spatial shifts in survey coverage of the strata in each region. Survey catchability *q*_*i,s*_ is therefore indexed by season and gear, *i*, and species, *s*. Moreover, the lower inflection points for the *q*_*i,s*_ priors were informed by the average survey coverage by gear and season. Survey coverage within each region was computed by dividing the average spatial coverage of strata across years by the total area of all strata (i.e., average area covered / area of survey domain). Survey coverage was then multiplied by 0.2 for deep-water species (Greenland Halibut, Atlantic Halibut, Witch Flounder, Redfish spp., White Hake, Silver Hake, and Monkfish) and 0.5 for the remainder. The lower range was widened, especially for deep-water species, to account for potential gear selectivity issues (e.g., escapement under the footgear; Walsh, 1992) and availability issues (e.g., portion of the stock in deeper water than covered by the survey). The upper inflection point was set to 1 as it is possible that the survey indices represent overestimates of the true population size in some instances.

A prior for observation error variance was informed by unbiased design-based estimates of survey variance associated with annual biomass estimates (Smith & Somerton, 1981). These estimates were used to calculate the coefficient of variation (CV) for each survey, year, and species; specifically,

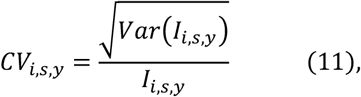

where *I* represents the design-based indices of biomass used in equation (2). These CVs were log transformed and survey, *i*, and species, *s*, specific means and standard deviations were used to inform the prior for 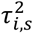 in log-space. Rather than treat the estimates of survey CV as a perfect indicator of total observation error, the prior was widened by multiplying the standard deviation of log CV by two to account for observation variance introduced by potential distributional shifts outside of the survey domain. Another two times multiplier was applied for deep-water species as they have more scope for shifting outside the survey domain.

#### Model selection

The degree to which density-dependence of the demersal fish community inhabiting the NL shelves is driven by intra versus interspecific competition is unknown. Nor is the degree to which species interactions affect population dynamics. It is, however, well known that the community has been fished for more than 500 years and this history was punctuated by a collapse of several stocks, most notably cod, in the early 1990s (Lear, 1998). This collapse was not isolated to commercial stocks, as the biomass of several non-commercial species collapsed at the same time. This was followed by a reorganization of the community, which implies that the system experienced a regime shift (Pedersen et al., 2017). Given this context, and the structure of the multispecies surplus production model described above, a series of hypotheses were tested:

##### Full

For this hypothesis, it is posited that population dynamics within regions are governed by an overarching system-level carrying capacity that affects all focal species as the aggregate biomass approaches system limits (i.e., inter-specific density-dependent effects are assumed). For this hypothesis and all others, annual reported landings of each species are accounted for in the production equation (5). Residual variations not explained by intrinsic growth, density-dependent effects or landings are described by process errors that are 1) correlated across species and said correlations are assumed to be unstructured, meaning relationships can be of differing strengths — positive, neutral, or negative — for each species-to-species pair; and 2) assumed to be temporally correlated, following an AR1 structure. Finally, a shift covariate was applied to the carrying capacity parameter to enable the estimation of different system limits before and after the community-wide collapse to assess support for a regime shift in each systems’ capacity for the focal demersal species.

##### Just shift

Model structure is the same as the **full** model, except the process errors are assumed to be independent and identically distributed random variables (*iid*) to assess the hypothesis that temporal and species correlations are explained by the shift.

##### Just correlation

Model structure is the same as the **full** model, except population dynamics are assumed to be affected by a common and time-invariant carrying capacity. The shift covariate was not applied.

##### Shared correlation

Model structure is the same as the **just correlation** model, except species by species correlations in the process errors are assumed to be the same across all pairs. This structure implies that there is a common but unknown environmental variable affecting the population dynamics of all species. The shift covariate was not applied.

##### Just species correlation

Model structure is the same as the **shared correlation** model, except the process errors at each time step are assumed to be *iid*. This structure implies that there is a common but unknown environmental process affecting all species, but the process is noisy with no temporal dependence. The shift covariate was not applied.

##### Just temporal correlation

Model structure is the same as the **shared correlation** model, except species by species correlations are assumed to be *iid*. This structure implies that environmental processes affect each species differently, however, there may be carry-over effects from one year to the next. The shift covariate was not applied.

##### No correlation

Model structure is the same as the **shared correlation** model, except both temporal and species correlations are assumed to be *iid*. That is, population dynamics are thought to be affected by a common time-invariant carrying capacity and residual variations not explained by intrinsic growth, inter-specific density-dependent effects or landings are noisy and independent across time and species. The shift covariate was not applied.

##### Single-species

Model structure is the same as the **no correlation** model, except population dynamics are assumed to be governed by species-specific carrying capacities (i.e., intra-specific density-dependent effects are assumed). This formulation is analogous to standard state-space Schaefer production models. The shift covariate was not applied.

The predictive ability of each of these models was tested using two cross-validation approaches: 1) leave-one-out cross-validation (LOO-CV), and 2) hindcast cross-validation (Hindcast-CV). LOO-CV is a form of exhaustive cross-validation where the model is repeatedly conditioned on a training set missing one observation (i.e., one fold) until the number of model folds equal the number of observations in the data. The missing observations are predicted at each fold, permitting assessments of the models’ ability to predict the actual value that was left-out at each fold. The hindcast-CV approach is similar, however it focuses on the models’ ability to predict the future. Under this approach, the model is repeatedly conditioned on a training set missing observations from the terminal year such that each fold excludes an increasing number of years worth of data from the tail of the time series (Kell et al., 2016). We folded back 20 years and, for each fold, predicted survey indices were compared to the observed survey indices (e.g., observed indices from 2020 were compared with predicted survey indices for 2020 from the model conditioned on data from 1978-2019). For both approaches, we denote predicted survey indices as 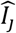 and the left out observations *I*_*j*_, where *j* represents all unique combinations of years, species, and survey indices present in the left out data. LOO-CV and Hindcast-CV prediction error scores for each model for each region were calculated by taking the mean squared error,

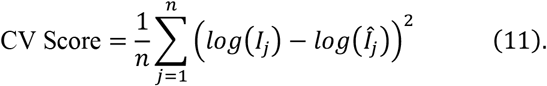

These scores were also averaged across methods (LOO-CV, Hindcast-CV) and region (Northeast NL Shelf, Grand Bank, Southern NL) to obtain an overall score of predictive ability of each model. A well-fitting model will result in predicted values that are close to the excluded values and, therefore, result in lower scores. Ultimately, these scores enable a model-comparison approach to hypothesis testing. These scores are comparable even for the single-species model as it compares species-specific data points to predictions.

### Code and Data Availability

All code used for the analyses in this study is publicly available in a research compendium on GitHub: https://github.com/PaulRegular/multispic. The compendium is organized as an R package, enabling straightforward replication of the results presented here and allowing the same methods to be applied to datasets from other regions.

## Results

Both cross-validation metrics (LOO-CV and Hindcast-CV scores) indicate that most multispecies production model formulations outperform a single-species production model when applied to seven species within three ecosystem production units (Northeast NL Shelf, Grand Bank, and Southern NL) off the east coast of Canada (Figure 2). Focusing on overall scores, the performance of the single-species production model was similar to a multispecies formulation that assumes there is no correlation in the process errors across species or time. There tends to be more notable decreases in the scores as temporal and species correlations are introduced, indicating an improvement in the predictive ability of these models. The “shared correlation” and “just correlation” formulations, in particular, tended to receive the lowest scores, and dropping the species and temporal correlations in lieu of a shift covariate (“just shift” formulation) resulted in a deterioration of predictive ability. Scores were improved when temporal and species-to-species correlations were introduced along with the shift covariate (“full” formulation); however, the fit of the “full” model tended to be poorer than the “just correlation” formulation, which further indicates that the “shift” covariate degraded the predictive ability of the model. Subsequent plots focus on the best fitting formulation, “just correlation”, to demonstrate model predictions.

**Fig 2:**
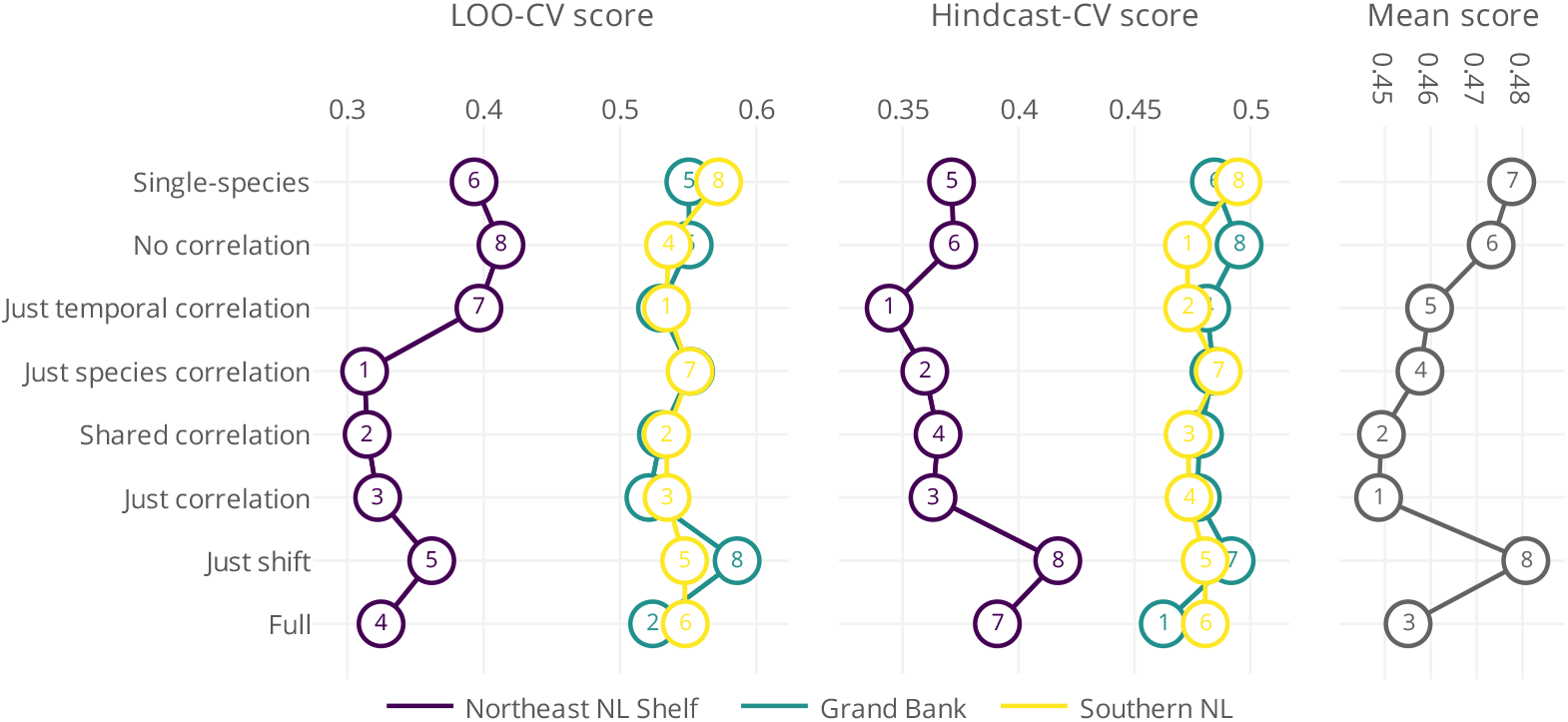
LOO-CV and Hindcast-CV scores from a series of production model formulations of increasing complexity, from a single-species production model (“Single-species”) to a multispecies production model with a covariate effect and species and temporal correlations (“Full”), fit to seven species in three ecosystem production units (Northeast NL Shelf, Grand Bank, and Southern NL) off the east coast of Canada. Overall scores for each model across score type and ecosystem production unit are shown on the rightmost facet.

The multispecies production model with unstructured species-to-species correlation and AR1 temporal correlation (“just correlation”) offered an explanation of the trends in survey indices of focal species across three ecosystem production units with little signs of systematic bias (see residual plots included in model dashboards; Supplement 1). Predicted indices track observed values and, by estimating catchability parameters by species and survey, indices from temporally fragmented surveys are stitched together and their trends are used to inform a continuous underlying trend in biomass (Figure 3). The earlier Yankee and Engel eras of the Canadian surveys tended to receive lower catchability estimates than the Campelen era survey; indices since 1996 therefore tend to be closer, in relative terms, to the underlying estimates of biomass from the model.

**Fig 3:**
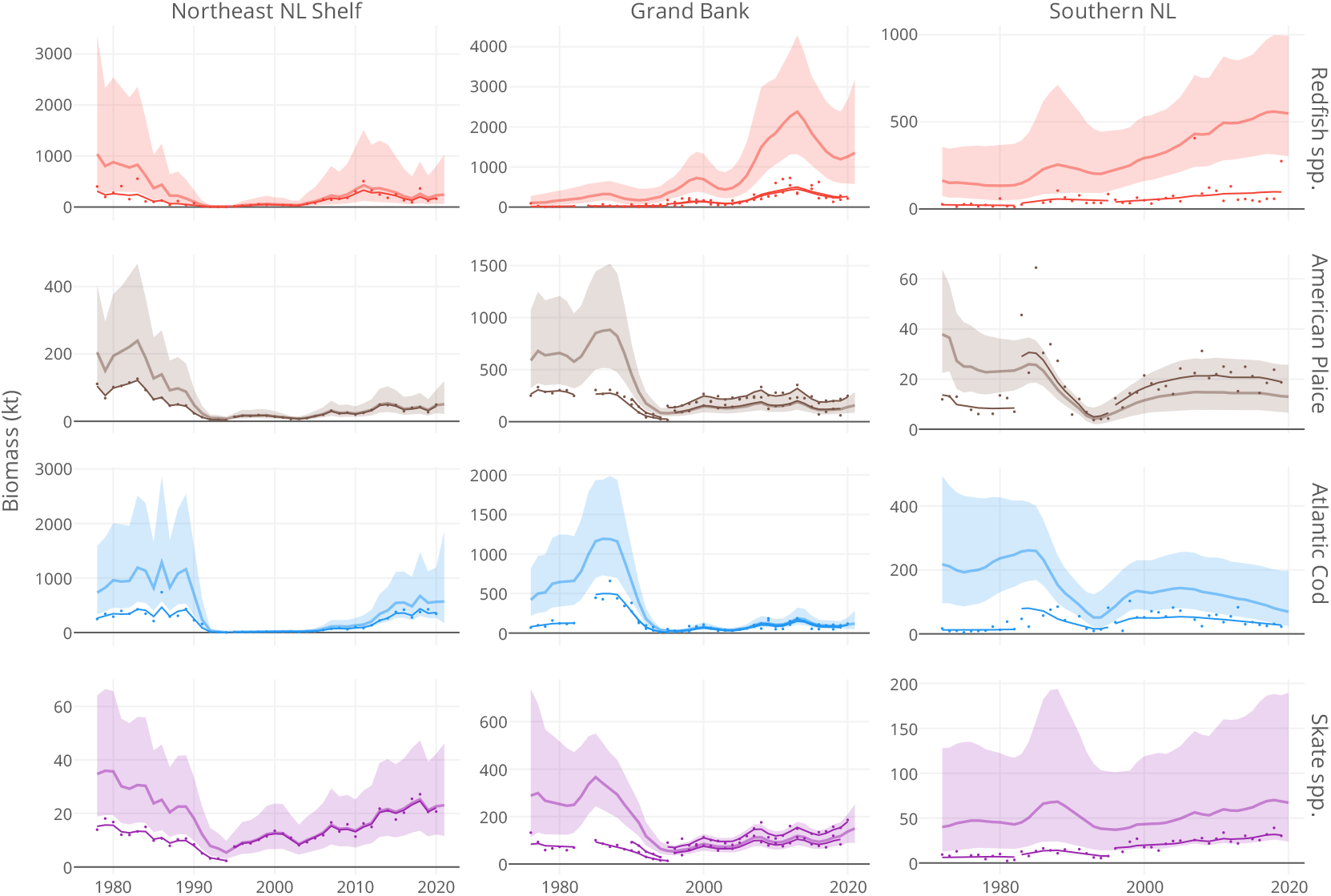
Trends in survey indices (dots) along with predicted indices (thin lines) and model estimates of biomass from the best fitting multispecies production model (thick lines). Uncertainty in the biomass estimates are represented by 95% confidence intervals (shaded region). Results from three ecosystem production units (Northeast NL Shelf, Grand Bank, and Southern NL) off the east coast of Canada are presented. Though data from seven species were used in each region, the species facet is limited to four for simplicity. Differences in biomass estimates from the indices stem from differing survey catchability estimates. Note differences in scale across facets.

Isolating residual changes in biomass not explained by reported fisheries landings or the production function (equation (4)) reveals substantive subtractions in the early 1990s across all three ecosystem production units (Figure 4). These residual subtractions exceed the absolute scale of landings taken in the late 1980s from the Northeast NL Shelf and Grand Bank ecosystem production units. The scale of changes derived from the production function and other processes are of a scale similar to landings reported in the Southern NL ecosystem production unit.

**Fig 4:**
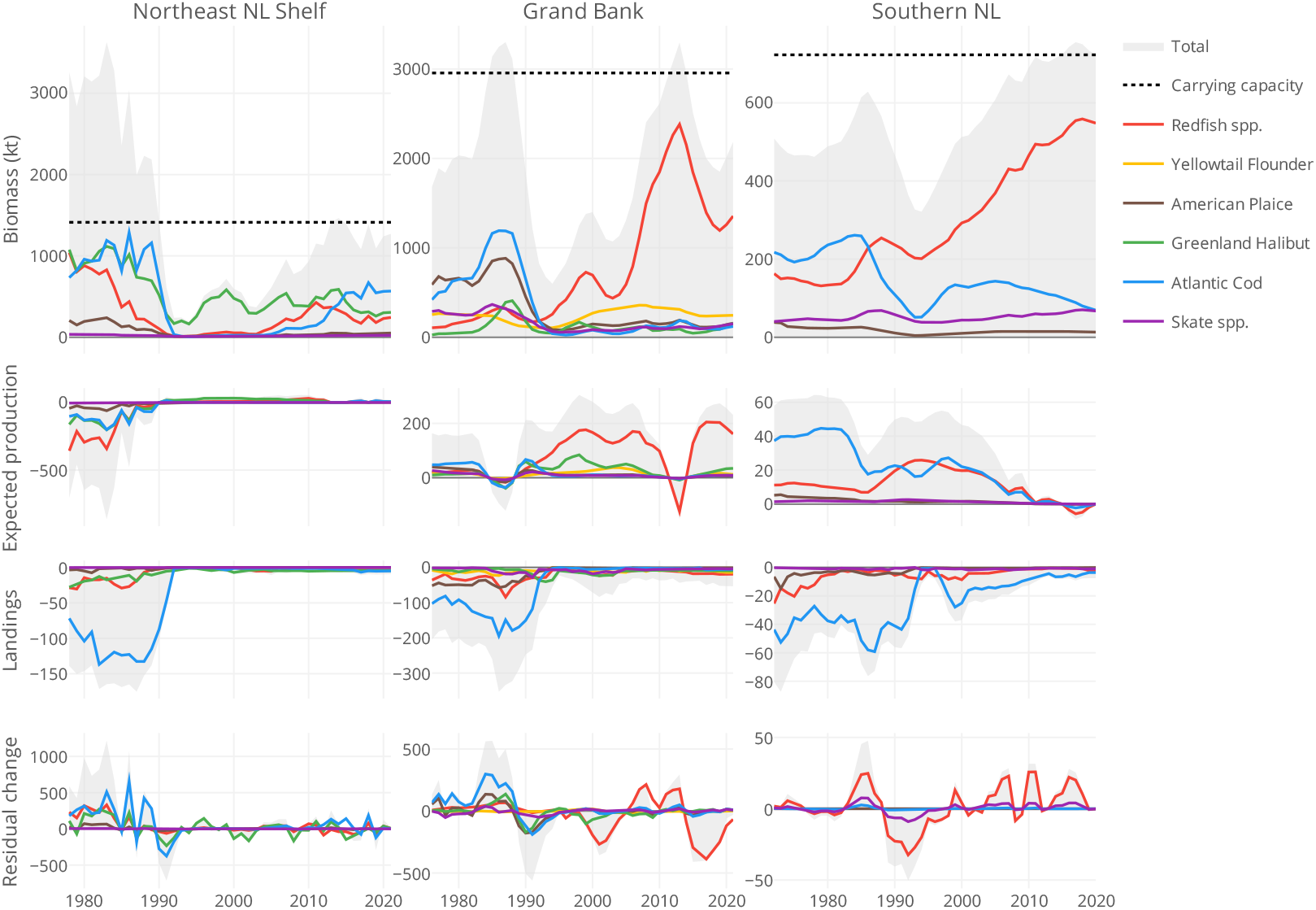
Trends in biomass, expected production, reported landings, and residual change from the best fitting multispecies production model across three ecosystem production units (Northeast NL Shelf, Grand Bank, and Southern NL) off the east coast of Canada. Species-specific trends from a subset of species are shown along with totals including seven species. Expected production represents expected changes from intrinsic growth and density-dependence (i.e., change from the production equation) and residual change represent process errors not explained by the production equation or reported landings. Note difference in scale across facets.

Estimates of biomass exceeded the carrying capacity estimated for the Northeast NL Shelf through the 1980s, and all focal species displayed abrupt declines in the early 1990s. Comparing community composition in the early 1980s to the 2010s, there are no clear shifts in the relative biomass of the focal species on the Northeast NL Shelf. Elsewhere, total biomass exceeded or approached system carrying capacity in the late 1980s, after which all species declined. Estimates of total biomass have gradually increased on the Grand Bank and off Southern NL since the mid 1990s, largely due to increasing Redfish spp. biomass estimates.

Further inspection of the process errors reveals common patterns across all focal species across three ecosystem production units (Figure 5). Like the exponentiated and unstandardized process errors (“residual change”) shown in Figure 4, the standardized process errors highlight substantive subtractions in the early 1990s, representing time-series lows for 19 out of 21 populations. The standardized values also reveal parallel and periodic increases and decreases within each region. The species-to-species correlations in the process errors estimated by the “just correlation” model indicate that 84% (53 of 63) of pairs were positively correlated. Also note that the “shared correlation” model estimates of a common species-to-species correlation parameter were 0.79 (95% CI: 0.70, 0.86), 0.68 (95% CI: 0.58, 0.76), and 0.81 (95% CI: 0.66, 0.90) for the Northeast NL Shelf, Grand Bank, and Southern NL ecosystem production units, respectively. This model received similar cross-validation scores as the “just correlation” model which, taken together, further supports the inference that the process errors are primarily positively correlated across species within each production unit.

**Fig 5:**
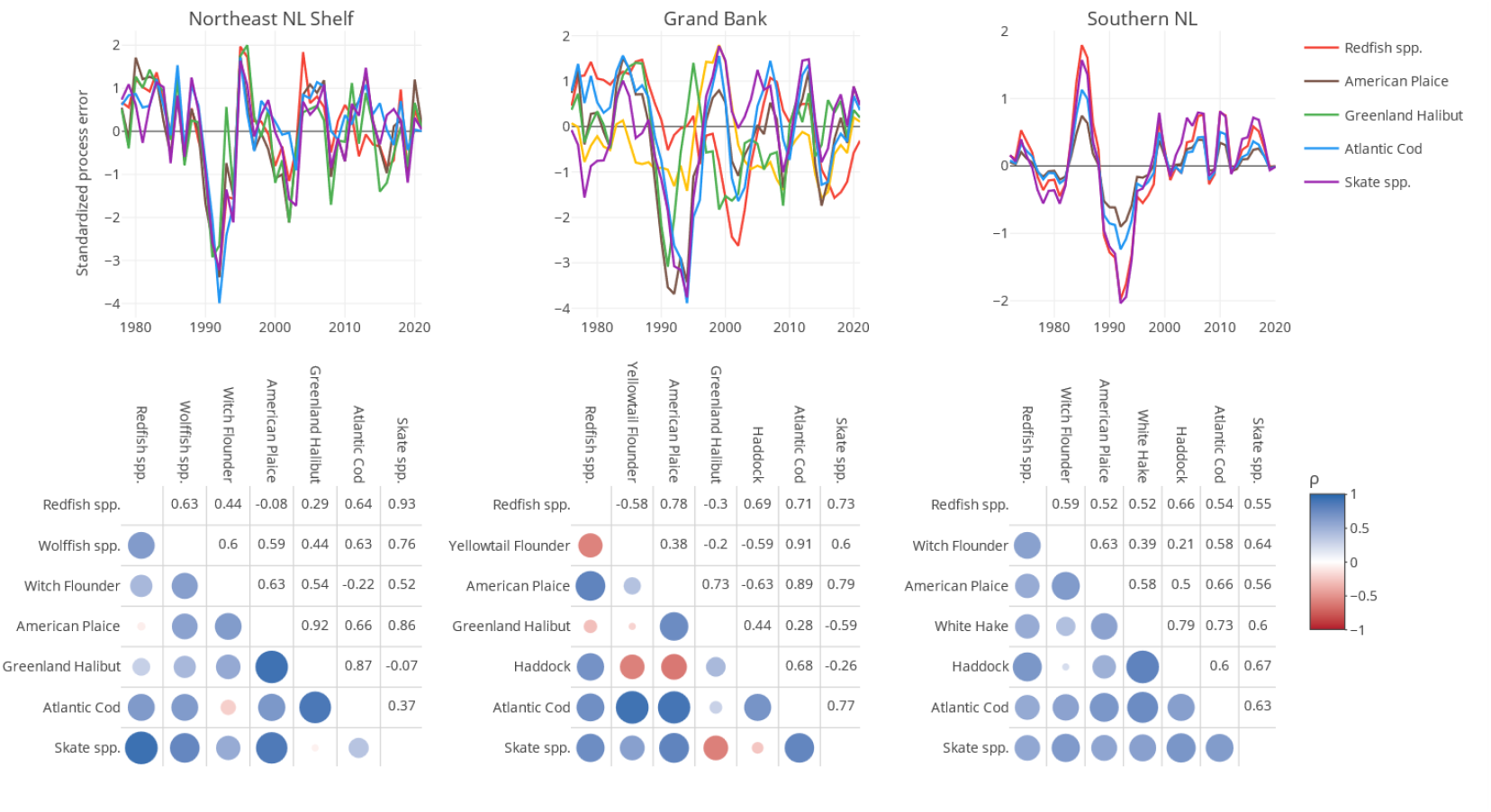
Trends (upper facet) and species-to-species correlations (lower facet) in standardized process errors from the best fitting multispecies production model applied to seven species within three ecosystem production units (Northeast NL Shelf, Grand Bank, and Southern NL) off the east coast of Canada. Process errors represent changes not explained by the production equation or reported landings. Points in the correlation plots are coloured and scaled by the level of species-to-species correlation, *ρ*.

## Discussion

Rather than assuming that populations are primarily regulated by species-specific carrying capacities, our multispecies production model assumes that both intra and interspecific competition stunts growth as total biomass approaches the environment’s maximal load. This assumption is conceptually similar to aggregate production models (Bundy et al., 2012; Fogarty et al., 2012; Mueter & Megrey, 2006) as it is rooted in the idea that total production, and consequently system-level MSY, is limited by the amount of resources available in a given ecosystem. In contrast to aggregate production models, we also attempt to capture the dynamics of species within a community. Fisheries landings, competitive interactions, predation, and prey availability all affect species-level production (Lotka, 1925; Schaefer, 1954; Volterra, 1926).

While our model explicitly accounts for landings, species interactions are implicitly accounted for by estimating species-to-species correlations (sensu Albertsen et al., 2018; see Gamble & Link, 2009 for a more explicit approach). Finally, by utilizing a state-space framework akin to single-species state-space production models (e.g., Millar & Meyer, 2000; Winker et al., 2018), we attempt to differentiate population processes from noise and bias from surveys of fish populations. The overall structure of the model allows species-specific dynamics to be captured while avoiding the assumption that the dynamics of each species is isolated and independent from other species sharing the same space and potentially competing for the same resources.

Our case study focuses on the population dynamics of commercially important demersal fish stocks off the east coast of Canada. Stocks in the area collapsed in the early 1990s (Lear, 1998), and the relative contribution of fishing and environmental impacts has been highly debated (Pedersen et al., 2017). We attempt to disentangle the impacts of fishing from environmental effects using our multispecies production model and, in doing so, we provide empirical evidence that environmental factors played a non-negligible role in the changes observed in the region.

First, we found general support for models with a system-level carrying capacity, consistent with the expectation that species within the same ecosystem production unit are constrained by a finite amount of available energy (Pepin et al., 2014). Second, we found evidence for synchronous changes in the demersal fish community, which implies that a common bottom-up driver is impacting the dynamics of these species (see also Bundy et al., 2012). This is supported by species-specific studies on capelin (*Mallotus villosus*) and Atlantic cod in the region which highlight the influence of bottom-up drivers (e.g., Buren, Koen-Alonso, & Stenson, 2014; Buren, Koen-Alonso, Pepin, et al., 2014; Koen-Alonso et al., 2021; Regular et al., 2022). Taken together, evidence is mounting that fishing was not the sole cause of the collapses observed in the early 1990s.

Our inference that environmental factors were a key driver of stock collapses was unexpected given the compelling narrative that fishing activity was the primary driver (e.g., Gomes et al., 1995; Hutchings, 1996). Since our model utilizes *reported* fisheries landings, a portion of these losses may be attributed to illegal fishing activity. However, it seems unlikely that the industry had the capacity to extract the amount needed to match the estimated losses. For instance, annual catches in the late 1980s across the Northeast NL Shelf and the Grand Bank totaled ∼450 kt while residual losses estimated by the model in the early 1990s was ∼1000 kt. The fishing industry would have had to covertly double its efforts to explain the declines. It follows that the decline must at least in part, if not primarily, be due to an unknown environmental driver. This contention is not new (Atkinson, 1994; see, for example, Morgan et al., 2002; Pedersen et al., 2017), however, it remains contentious and perplexing as we lack specific causal explanations.

While increasingly cold conditions through the 1980s and early 1990s undoubtedly affected the distribution of multiple species Robertson et al. (2021), it is not yet clear whether shifting temperatures was the primary driver of the collapse and, if it was, the causal pathway has yet to be determined.

Regardless of the environmental driver behind the 1990s collapse, it is possible that increasingly industrialized and intense fishing activity through the 1960s and 1970s reduced population diversity and, consequently, hampered the ability of the species within the community to buffer subsequent environmental changes (*sensu* the portfolio effect, Schindler et al., 2010). Yet, a recent study found no evidence of genetic diversity loss in heavily exploited species like Atlantic cod (Pinsky et al., 2021). Another hypothesis is that fishing activity bounded the safe operating space of the system, triggering an alternate stable state (Scheffer et al., 2015). While not a perfect test of chaotic dynamics, we did assess the possibility of a systematic shift in system-level carrying capacities and found little support for this hypothesis. That said, there were clear shifts in the communities in the region and these shifts may have emerged from the combined effects of interspecific competition and shifting energy pathways. It is well known that the dominant forage species in the area shifted from capelin to shrimp (Dawe et al., 2012) and this change was detrimental for cod (Link & Sherwood, 2019; Mullowney & Rose, 2014; Regular et al., 2022) and perhaps other piscivorous species that rely on capelin. Shrimp are an important prey item for redfish species (Brown-Vuillemin et al., 2022), so it is possible that the increasing shrimp population helped support concurrent recruitment pulses of redfish. We admit that this conjecture is highly speculative; however, we add it as a simple example of how bottom-up forces may be driving the observed changes in the community. The reality is obviously more complex and the observed restructuring of the communities may be akin to the “paradox of plankton” where the continuous interaction of ecological and environmental factors give rise to “oscillations and chaos, with a continuous wax and wane of species within the community” (Scheffer et al., 2003).

Like all models, our multispecies production model is an imperfect abstraction of nature and while it may be useful in some contexts, it is important to consider its limitations when interpreting results. First, it is important to remember that there may be a spatial mismatch in the structure and function of the populations included in this study as some stock boundaries differ from the regions used in this study. For instance, Atlantic cod in NAFO divisions 2J3KL are considered a separate stock from cod in divisions 3NO (Templeman, 1962) and here we split 2J3K (Northeast NL Shelf) and 3LNO (Grand Bank) into distinct regions. Assuming that our results are comparable to previous results, it is peculiar that they indicate that total biomass in the Northeast NL Shelf production unit was above the carrying capacity of that region through the 1980s. This finding contradicts historic records that suggest populations such as Atlantic cod in 2J3KL were at substantially higher levels in the 1970s and earlier (Rose, 2004; Schijns et al., 2021), which implies that the carrying capacity should be higher than estimated by the model presented here. Still, it is possible that the 1970s represents a period of unusually high productivity, where the system may have exceeded the carrying capacity.

Results from the Grand Bank and Southern NL also indicate that the demersal fish community is currently dominated by redfish and, consequently, the system appears to be approaching its carrying capacity. Though redfish are currently rebounding in parts of eastern Canada (Cadigan et al., 2022), the implication that it dominates the benthic community seems unrealistic. This result may be an artifact of low estimates of survey catchability or the model’s inability to properly account for year effects. Observation errors are assumed to be lognormally distributed, however, extreme catch events / black swan events in space can introduce ‘year effects’ that may be better accounted for by assuming a distribution with heavier tails, such as the t-distribution (Anderson & Ward, 2019).

## Conclusion

Practitioners are becoming increasingly aware of the need to apply an ecosystem based fisheries management (EBFM; Pikitch et al., 2004). A robust understanding of the interactions of multiple species with each other and their environment is a critical prerequisite for advancing a EBFM. There are multiple analytical pathways to support such management, however, data requirements are often prohibitive (Latour et al., 2003). Our study documents a production model that can estimate the population dynamics of multiple stocks using commonly available landings and survey data. This approach enables the estimation of multispecies trends and provides an avenue for producing species-specific projections conditioned on recent community dynamics. As such, it may serve as a relatively tractable method for informing management decisions for multiple species occupying the same region. While our model has limitations, it represents a step forward in understanding the complex interactions among species in marine ecosystems and provides a framework for supporting sustainable management decisions.

## Acknowledgements

We thank the many ship’s crew and research staff involved in leading the surveys and collecting the data used in this study. These surveys were supported by Fisheries and Oceans Canada (DFO). An earlier draft version of the abstract, introduction, and discussion sections were written with the assistance of ChatGPT (March 14, 2023 version).

