## Supplement 1 for "Summing the parts: Improving population estimates using a state-space multispecies production model": Just correlation - Grand Bank.html

multispic diagnostics and results


multispic diagnostics and results

### Inputs

#### Row

##### Index

##### Landings

##### Covariates

N/A; no process error or K covariates supplied

### Residuals

#### Row

##### Residuals ~ predicted value

##### ~ year

##### ~ survey

##### ~ species

### Parameters

#### Row

##### Correlation

##### log(K)

##### log(r)

##### log(B0)

##### log(SD_B_)

##### logit(rho)

##### logit(phi)

##### log(q)

##### log(SD_I_)

##### beta_pe_

N/A; no process error covariates supplied

##### beta_K_

N/A; no K covariates supplied

##### Estimates (plot)

##### Estimates (table)

| parameter | group | estimate | CV | lower | upper |
| --- | --- | --- | --- | --- | --- |
| K | All species | 2955.346 | 1.176 | 2149.412 | 4063.468 |
| r | American Plaice-3LNO | 0.155 | 1.724 | 0.053 | 0.450 |
|  | Atlantic Cod-3LNO | 0.262 | 1.795 | 0.083 | 0.826 |
|  | Greenland Halibut-3LNO | 0.864 | 1.459 | 0.412 | 1.812 |
|  | Haddock-3LNO | 0.508 | 1.512 | 0.226 | 1.141 |
|  | Redfish spp.-3LNO | 0.455 | 1.506 | 0.204 | 1.016 |
|  | Skate spp.-3LNO | 0.210 | 1.488 | 0.096 | 0.458 |
|  | Yellowtail Flounder-3LNO | 0.184 | 1.332 | 0.105 | 0.323 |
| B0 | American Plaice-3LNO | 533.020 | 1.408 | 272.444 | 1042.824 |
|  | Atlantic Cod-3LNO | 351.102 | 1.556 | 147.560 | 835.406 |
|  | Greenland Halibut-3LNO | 22.756 | 2.197 | 4.866 | 106.426 |
|  | Haddock-3LNO | 1.271 | 2.216 | 0.267 | 6.041 |
|  | Redfish spp.-3LNO | 97.980 | 1.760 | 32.370 | 296.569 |
|  | Skate spp.-3LNO | 290.874 | 1.696 | 103.290 | 819.130 |
|  | Yellowtail Flounder-3LNO | 251.716 | 1.375 | 134.866 | 469.804 |
| sd\_B | American Plaice-3LNO | 0.133 | 1.207 | 0.092 | 0.192 |
|  | Atlantic Cod-3LNO | 0.228 | 1.204 | 0.158 | 0.328 |
|  | Greenland Halibut-3LNO | 0.310 | 1.225 | 0.208 | 0.462 |
|  | Haddock-3LNO | 0.209 | 1.316 | 0.122 | 0.358 |
|  | Redfish spp.-3LNO | 0.157 | 1.318 | 0.091 | 0.269 |
|  | Skate spp.-3LNO | 0.116 | 1.213 | 0.079 | 0.169 |
|  | Yellowtail Flounder-3LNO | 0.045 | 1.577 | 0.018 | 0.109 |
| rho | Atlantic Cod-3LNO - American Plaice-3LNO | 0.893 | 0.618 | 0.023 | 0.993 |
|  | Greenland Halibut-3LNO - American Plaice-3LNO | 0.733 | 0.566 | -0.311 | 0.975 |
|  | Haddock-3LNO - American Plaice-3LNO | -0.628 | 0.550 | -0.960 | 0.441 |
|  | Redfish spp.-3LNO - American Plaice-3LNO | 0.785 | 0.646 | -0.421 | 0.988 |
|  | Skate spp.-3LNO - American Plaice-3LNO | 0.787 | 0.655 | -0.442 | 0.989 |
|  | Yellowtail Flounder-3LNO - American Plaice-3LNO | 0.379 | 0.465 | -0.529 | 0.882 |
|  | Greenland Halibut-3LNO - Atlantic Cod-3LNO | 0.282 | 0.518 | -0.683 | 0.888 |
|  | Haddock-3LNO - Atlantic Cod-3LNO | 0.680 | 0.690 | -0.682 | 0.986 |
|  | Redfish spp.-3LNO - Atlantic Cod-3LNO | 0.713 | 0.650 | -0.555 | 0.984 |
|  | Skate spp.-3LNO - Atlantic Cod-3LNO | 0.772 | 0.657 | -0.476 | 0.988 |
|  | Yellowtail Flounder-3LNO - Atlantic Cod-3LNO | 0.912 | 0.622 | 0.114 | 0.995 |
|  | Haddock-3LNO - Greenland Halibut-3LNO | 0.439 | 0.503 | -0.547 | 0.915 |
|  | Redfish spp.-3LNO - Greenland Halibut-3LNO | -0.295 | 0.428 | -0.834 | 0.532 |
|  | Skate spp.-3LNO - Greenland Halibut-3LNO | -0.593 | 0.572 | -0.961 | 0.533 |
|  | Yellowtail Flounder-3LNO - Greenland Halibut-3LNO | -0.204 | 0.488 | -0.849 | 0.686 |
|  | Redfish spp.-3LNO - Haddock-3LNO | 0.692 | 0.674 | -0.637 | 0.985 |
|  | Skate spp.-3LNO - Haddock-3LNO | -0.264 | 0.705 | -0.963 | 0.896 |
|  | Yellowtail Flounder-3LNO - Haddock-3LNO | -0.591 | 0.712 | -0.984 | 0.788 |
|  | Skate spp.-3LNO - Redfish spp.-3LNO | 0.726 | 0.698 | -0.647 | 0.989 |
|  | Yellowtail Flounder-3LNO - Redfish spp.-3LNO | -0.579 | 0.731 | -0.986 | 0.822 |
|  | Yellowtail Flounder-3LNO - Skate spp.-3LNO | 0.598 | 0.771 | -0.865 | 0.991 |
| phi | All species | 0.538 | 0.602 | 0.342 | 0.723 |
| q | American Plaice-3LNO-Fall-Campelen | 1.676 | 1.269 | 1.051 | 2.673 |
|  | American Plaice-3LNO-Fall-Engel | 0.497 | 1.273 | 0.310 | 0.798 |
|  | American Plaice-3LNO-Spring-Campelen | 1.052 | 1.273 | 0.656 | 1.689 |
|  | American Plaice-3LNO-Spring-Engel | 0.311 | 1.274 | 0.193 | 0.500 |
|  | American Plaice-3LNO-Spring-Yankee | 0.450 | 1.353 | 0.249 | 0.814 |
|  | Atlantic Cod-3LNO-Fall-Campelen | 0.952 | 1.264 | 0.602 | 1.507 |
|  | Atlantic Cod-3LNO-Fall-Engel | 0.410 | 1.282 | 0.252 | 0.666 |
|  | Atlantic Cod-3LNO-Spring-Campelen | 0.809 | 1.273 | 0.504 | 1.297 |
|  | Atlantic Cod-3LNO-Spring-Engel | 0.419 | 1.250 | 0.271 | 0.649 |
|  | Atlantic Cod-3LNO-Spring-Yankee | 0.191 | 1.370 | 0.103 | 0.354 |
|  | Greenland Halibut-3LNO-Fall-Campelen | 0.230 | 1.398 | 0.119 | 0.443 |
|  | Greenland Halibut-3LNO-Fall-Engel | 0.090 | 1.317 | 0.052 | 0.154 |
|  | Greenland Halibut-3LNO-Spring-Campelen | 0.168 | 1.408 | 0.086 | 0.328 |
|  | Greenland Halibut-3LNO-Spring-Engel | 0.023 | 1.308 | 0.014 | 0.039 |
|  | Greenland Halibut-3LNO-Spring-Yankee | 0.070 | 1.910 | 0.020 | 0.248 |
|  | Haddock-3LNO-Fall-Campelen | 0.565 | 1.337 | 0.319 | 0.999 |
|  | Haddock-3LNO-Fall-Engel | 0.371 | 1.385 | 0.196 | 0.703 |
|  | Haddock-3LNO-Spring-Campelen | 1.114 | 1.348 | 0.620 | 1.999 |
|  | Haddock-3LNO-Spring-Engel | 0.667 | 1.299 | 0.399 | 1.112 |
|  | Haddock-3LNO-Spring-Yankee | 0.307 | 1.628 | 0.118 | 0.797 |
|  | Redfish spp.-3LNO-Fall-Campelen | 0.210 | 1.338 | 0.119 | 0.371 |
|  | Redfish spp.-3LNO-Fall-Engel | 0.139 | 1.633 | 0.053 | 0.365 |
|  | Redfish spp.-3LNO-Spring-Campelen | 0.189 | 1.342 | 0.106 | 0.336 |
|  | Redfish spp.-3LNO-Spring-Engel | 0.110 | 1.465 | 0.052 | 0.233 |
|  | Redfish spp.-3LNO-Spring-Yankee | 0.124 | 1.784 | 0.040 | 0.385 |
|  | Skate spp.-3LNO-Fall-Campelen | 1.305 | 1.215 | 0.891 | 1.911 |
|  | Skate spp.-3LNO-Fall-Engel | 0.446 | 1.220 | 0.302 | 0.658 |
|  | Skate spp.-3LNO-Spring-Campelen | 0.882 | 1.217 | 0.600 | 1.297 |
|  | Skate spp.-3LNO-Spring-Engel | 0.254 | 1.216 | 0.173 | 0.372 |
|  | Skate spp.-3LNO-Spring-Yankee | 0.289 | 1.441 | 0.141 | 0.591 |
|  | Yellowtail Flounder-3LNO-Fall-Campelen | 1.106 | 1.271 | 0.691 | 1.768 |
|  | Yellowtail Flounder-3LNO-Fall-Engel | 0.424 | 1.256 | 0.271 | 0.663 |
|  | Yellowtail Flounder-3LNO-Spring-Campelen | 1.080 | 1.275 | 0.671 | 1.739 |
|  | Yellowtail Flounder-3LNO-Spring-Engel | 0.432 | 1.230 | 0.288 | 0.648 |
|  | Yellowtail Flounder-3LNO-Spring-Yankee | 0.391 | 1.313 | 0.229 | 0.667 |
| sd\_I | American Plaice-3LNO-Fall-Campelen | 0.122 | 1.229 | 0.081 | 0.182 |
|  | American Plaice-3LNO-Fall-Engel | 0.096 | 1.228 | 0.064 | 0.144 |
|  | American Plaice-3LNO-Spring-Campelen | 0.264 | 1.173 | 0.193 | 0.360 |
|  | American Plaice-3LNO-Spring-Engel | 0.202 | 1.370 | 0.109 | 0.374 |
|  | American Plaice-3LNO-Spring-Yankee | 0.080 | 1.448 | 0.039 | 0.165 |
|  | Atlantic Cod-3LNO-Fall-Campelen | 0.259 | 1.187 | 0.185 | 0.363 |
|  | Atlantic Cod-3LNO-Fall-Engel | 0.194 | 1.357 | 0.106 | 0.352 |
|  | Atlantic Cod-3LNO-Spring-Campelen | 0.447 | 1.179 | 0.324 | 0.617 |
|  | Atlantic Cod-3LNO-Spring-Engel | 0.321 | 1.265 | 0.202 | 0.509 |
|  | Atlantic Cod-3LNO-Spring-Yankee | 0.174 | 1.406 | 0.089 | 0.340 |
|  | Greenland Halibut-3LNO-Fall-Campelen | 0.137 | 1.345 | 0.077 | 0.245 |
|  | Greenland Halibut-3LNO-Fall-Engel | 0.150 | 1.553 | 0.063 | 0.356 |
|  | Greenland Halibut-3LNO-Spring-Campelen | 0.355 | 1.180 | 0.257 | 0.491 |
|  | Greenland Halibut-3LNO-Spring-Engel | 0.207 | 1.528 | 0.090 | 0.475 |
|  | Greenland Halibut-3LNO-Spring-Yankee | 0.420 | 1.372 | 0.226 | 0.779 |
|  | Haddock-3LNO-Fall-Campelen | 0.846 | 1.158 | 0.634 | 1.127 |
|  | Haddock-3LNO-Fall-Engel | 0.560 | 1.421 | 0.281 | 1.116 |
|  | Haddock-3LNO-Spring-Campelen | 0.598 | 1.178 | 0.434 | 0.824 |
|  | Haddock-3LNO-Spring-Engel | 0.772 | 1.221 | 0.522 | 1.142 |
|  | Haddock-3LNO-Spring-Yankee | 1.103 | 1.335 | 0.626 | 1.941 |
|  | Redfish spp.-3LNO-Fall-Campelen | 0.401 | 1.181 | 0.289 | 0.556 |
|  | Redfish spp.-3LNO-Fall-Engel | 0.757 | 1.380 | 0.402 | 1.424 |
|  | Redfish spp.-3LNO-Spring-Campelen | 0.485 | 1.174 | 0.354 | 0.664 |
|  | Redfish spp.-3LNO-Spring-Engel | 0.888 | 1.246 | 0.577 | 1.366 |
|  | Redfish spp.-3LNO-Spring-Yankee | 1.202 | 1.355 | 0.663 | 2.179 |
|  | Skate spp.-3LNO-Fall-Campelen | 0.145 | 1.201 | 0.101 | 0.208 |
|  | Skate spp.-3LNO-Fall-Engel | 0.072 | 1.449 | 0.035 | 0.149 |
|  | Skate spp.-3LNO-Spring-Campelen | 0.162 | 1.173 | 0.118 | 0.221 |
|  | Skate spp.-3LNO-Spring-Engel | 0.094 | 1.422 | 0.047 | 0.187 |
|  | Skate spp.-3LNO-Spring-Yankee | 0.232 | 1.540 | 0.100 | 0.541 |
|  | Yellowtail Flounder-3LNO-Fall-Campelen | 0.188 | 1.171 | 0.138 | 0.256 |
|  | Yellowtail Flounder-3LNO-Fall-Engel | 0.230 | 1.238 | 0.152 | 0.350 |
|  | Yellowtail Flounder-3LNO-Spring-Campelen | 0.240 | 1.149 | 0.183 | 0.315 |
|  | Yellowtail Flounder-3LNO-Spring-Engel | 0.184 | 1.237 | 0.121 | 0.279 |
|  | Yellowtail Flounder-3LNO-Spring-Yankee | 0.173 | 1.292 | 0.105 | 0.285 |

### Population trends

#### Row

##### Observed and predicted index

##### Process error

##### Standardized process error

> Standardized process error = process error, in log space, divided by
> the standard deviation of the process.

##### Process error correlation

##### Biomass

##### Total biomass

##### Harvest rate
