## Supplement 1 for "Summing the parts: Improving population estimates using a state-space multispecies production model": Just correlation - Northeast NL Shelf.html

multispic diagnostics and results


multispic diagnostics and results

### Inputs

#### Row

##### Index

##### Landings

##### Covariates

N/A; no process error or K covariates supplied

### Residuals

#### Row

##### Residuals ~ predicted value

##### ~ year

##### ~ survey

##### ~ species

### Parameters

#### Row

##### Correlation

##### log(K)

##### log(r)

##### log(B0)

##### log(SD_B_)

##### logit(rho)

##### logit(phi)

##### log(q)

##### log(SD_I_)

##### beta_pe_

N/A; no process error covariates supplied

##### beta_K_

N/A; no K covariates supplied

##### Estimates (plot)

##### Estimates (table)

| parameter | group | estimate | CV | lower | upper |
| --- | --- | --- | --- | --- | --- |
| K | All species | 1412.663 | 1.339 | 797.169 | 2503.382 |
| r | American Plaice-2J3K | 0.162 | 1.431 | 0.080 | 0.326 |
|  | Atlantic Cod-2J3K | 0.107 | 1.909 | 0.030 | 0.382 |
|  | Greenland Halibut-2J3K | 0.115 | 1.540 | 0.049 | 0.269 |
|  | Redfish spp.-2J3K | 0.263 | 1.421 | 0.132 | 0.524 |
|  | Skate spp.-2J3K | 0.090 | 1.442 | 0.044 | 0.185 |
|  | Witch Flounder-2J3K | 0.143 | 1.572 | 0.059 | 0.348 |
|  | Wolffish spp.-2J3K | 0.182 | 1.430 | 0.090 | 0.367 |
| B0 | American Plaice-2J3K | 176.908 | 1.538 | 76.088 | 411.320 |
|  | Atlantic Cod-2J3K | 552.439 | 1.806 | 173.347 | 1760.565 |
|  | Greenland Halibut-2J3K | 966.881 | 1.794 | 307.414 | 3041.039 |
|  | Redfish spp.-2J3K | 809.136 | 1.974 | 213.300 | 3069.390 |
|  | Skate spp.-2J3K | 31.078 | 1.415 | 15.745 | 61.341 |
|  | Witch Flounder-2J3K | 77.325 | 1.657 | 28.731 | 208.107 |
|  | Wolffish spp.-2J3K | 55.726 | 1.587 | 22.540 | 137.768 |
| sd\_B | American Plaice-2J3K | 0.267 | 1.139 | 0.207 | 0.345 |
|  | Atlantic Cod-2J3K | 0.462 | 1.181 | 0.334 | 0.640 |
|  | Greenland Halibut-2J3K | 0.208 | 1.176 | 0.152 | 0.287 |
|  | Redfish spp.-2J3K | 0.372 | 1.215 | 0.254 | 0.545 |
|  | Skate spp.-2J3K | 0.151 | 1.208 | 0.104 | 0.218 |
|  | Witch Flounder-2J3K | 0.317 | 1.157 | 0.238 | 0.422 |
|  | Wolffish spp.-2J3K | 0.316 | 1.146 | 0.242 | 0.412 |
| rho | Atlantic Cod-2J3K - American Plaice-2J3K | 0.656 | 0.435 | -0.126 | 0.935 |
|  | Greenland Halibut-2J3K - American Plaice-2J3K | 0.918 | 0.602 | 0.209 | 0.994 |
|  | Redfish spp.-2J3K - American Plaice-2J3K | -0.081 | 0.253 | -0.528 | 0.401 |
|  | Skate spp.-2J3K - American Plaice-2J3K | 0.857 | 0.622 | -0.143 | 0.991 |
|  | Witch Flounder-2J3K - American Plaice-2J3K | 0.628 | 0.520 | -0.373 | 0.953 |
|  | Wolffish spp.-2J3K - American Plaice-2J3K | 0.586 | 0.537 | -0.465 | 0.952 |
|  | Greenland Halibut-2J3K - Atlantic Cod-2J3K | 0.873 | 0.616 | -0.061 | 0.992 |
|  | Redfish spp.-2J3K - Atlantic Cod-2J3K | 0.640 | 0.545 | -0.414 | 0.961 |
|  | Skate spp.-2J3K - Atlantic Cod-2J3K | 0.367 | 0.421 | -0.459 | 0.852 |
|  | Witch Flounder-2J3K - Atlantic Cod-2J3K | -0.224 | 0.433 | -0.813 | 0.592 |
|  | Wolffish spp.-2J3K - Atlantic Cod-2J3K | 0.635 | 0.463 | -0.229 | 0.939 |
|  | Redfish spp.-2J3K - Greenland Halibut-2J3K | 0.291 | 0.323 | -0.342 | 0.742 |
|  | Skate spp.-2J3K - Greenland Halibut-2J3K | -0.067 | 0.340 | -0.641 | 0.556 |
|  | Witch Flounder-2J3K - Greenland Halibut-2J3K | 0.536 | 0.578 | -0.600 | 0.955 |
|  | Wolffish spp.-2J3K - Greenland Halibut-2J3K | 0.440 | 0.480 | -0.502 | 0.905 |
|  | Skate spp.-2J3K - Redfish spp.-2J3K | 0.926 | 0.605 | 0.250 | 0.995 |
|  | Witch Flounder-2J3K - Redfish spp.-2J3K | 0.443 | 0.486 | -0.511 | 0.908 |
|  | Wolffish spp.-2J3K - Redfish spp.-2J3K | 0.630 | 0.597 | -0.543 | 0.970 |
|  | Witch Flounder-2J3K - Skate spp.-2J3K | 0.524 | 0.602 | -0.654 | 0.960 |
|  | Wolffish spp.-2J3K - Skate spp.-2J3K | 0.765 | 0.634 | -0.430 | 0.986 |
|  | Wolffish spp.-2J3K - Witch Flounder-2J3K | 0.604 | 0.518 | -0.401 | 0.949 |
| phi | All species | 0.130 | 0.690 | 0.030 | 0.419 |
| q | American Plaice-2J3K-Fall-Campelen | 0.941 | 1.383 | 0.498 | 1.778 |
|  | American Plaice-2J3K-Fall-Engel | 0.509 | 1.394 | 0.265 | 0.975 |
|  | Atlantic Cod-2J3K-Fall-Campelen | 0.649 | 1.398 | 0.336 | 1.251 |
|  | Atlantic Cod-2J3K-Fall-Engel | 0.360 | 1.377 | 0.192 | 0.673 |
|  | Greenland Halibut-2J3K-Fall-Campelen | 0.427 | 1.848 | 0.128 | 1.422 |
|  | Greenland Halibut-2J3K-Fall-Engel | 0.157 | 1.731 | 0.054 | 0.461 |
|  | Redfish spp.-2J3K-Fall-Campelen | 0.777 | 1.827 | 0.238 | 2.532 |
|  | Redfish spp.-2J3K-Fall-Engel | 0.298 | 1.628 | 0.115 | 0.773 |
|  | Skate spp.-2J3K-Fall-Campelen | 0.973 | 1.360 | 0.533 | 1.778 |
|  | Skate spp.-2J3K-Fall-Engel | 0.437 | 1.352 | 0.242 | 0.788 |
|  | Witch Flounder-2J3K-Fall-Campelen | 0.350 | 1.588 | 0.141 | 0.867 |
|  | Witch Flounder-2J3K-Fall-Engel | 0.225 | 1.444 | 0.109 | 0.462 |
|  | Wolffish spp.-2J3K-Fall-Campelen | 0.729 | 1.392 | 0.381 | 1.392 |
|  | Wolffish spp.-2J3K-Fall-Engel | 0.676 | 1.407 | 0.346 | 1.319 |
| sd\_I | American Plaice-2J3K-Fall-Campelen | 0.113 | 1.390 | 0.059 | 0.215 |
|  | American Plaice-2J3K-Fall-Engel | 0.088 | 1.519 | 0.039 | 0.199 |
|  | Atlantic Cod-2J3K-Fall-Campelen | 0.179 | 1.548 | 0.076 | 0.421 |
|  | Atlantic Cod-2J3K-Fall-Engel | 0.313 | 1.372 | 0.168 | 0.581 |
|  | Greenland Halibut-2J3K-Fall-Campelen | 0.071 | 1.566 | 0.029 | 0.170 |
|  | Greenland Halibut-2J3K-Fall-Engel | 0.111 | 1.422 | 0.056 | 0.221 |
|  | Redfish spp.-2J3K-Fall-Campelen | 0.274 | 1.300 | 0.164 | 0.458 |
|  | Redfish spp.-2J3K-Fall-Engel | 0.505 | 1.261 | 0.321 | 0.795 |
|  | Skate spp.-2J3K-Fall-Campelen | 0.097 | 1.270 | 0.061 | 0.155 |
|  | Skate spp.-2J3K-Fall-Engel | 0.100 | 1.324 | 0.058 | 0.174 |
|  | Witch Flounder-2J3K-Fall-Campelen | 0.088 | 1.948 | 0.024 | 0.324 |
|  | Witch Flounder-2J3K-Fall-Engel | 0.119 | 1.824 | 0.037 | 0.388 |
|  | Wolffish spp.-2J3K-Fall-Campelen | 0.134 | 1.372 | 0.072 | 0.249 |
|  | Wolffish spp.-2J3K-Fall-Engel | 0.066 | 1.968 | 0.018 | 0.250 |

### Population trends

#### Row

##### Observed and predicted index

##### Process error

##### Standardized process error

> Standardized process error = process error, in log space, divided by
> the standard deviation of the process.

##### Process error correlation

##### Biomass

##### Total biomass

##### Harvest rate
