## Supplement 1 for "Summing the parts: Improving population estimates using a state-space multispecies production model": Just correlation - Southern NL.html

multispic diagnostics and results


multispic diagnostics and results

### Inputs

#### Row

##### Index

##### Landings

##### Covariates

N/A; no process error or K covariates supplied

### Residuals

#### Row

##### Residuals ~ predicted value

##### ~ year

##### ~ survey

##### ~ species

### Parameters

#### Row

##### Correlation

##### log(K)

##### log(r)

##### log(B0)

##### log(SD_B_)

##### logit(rho)

##### logit(phi)

##### log(q)

##### log(SD_I_)

##### beta_pe_

N/A; no process error covariates supplied

##### beta_K_

N/A; no K covariates supplied

##### Estimates (plot)

##### Estimates (table)

| parameter | group | estimate | CV | lower | upper |
| --- | --- | --- | --- | --- | --- |
| K | All species | 723.111 | 1.222 | 488.397 | 1070.625 |
| r | American Plaice-3Ps | 0.461 | 1.315 | 0.269 | 0.788 |
|  | Atlantic Cod-3Ps | 0.579 | 1.379 | 0.308 | 1.087 |
|  | Haddock-3Ps | 0.271 | 1.559 | 0.114 | 0.647 |
|  | Redfish spp.-3Ps | 0.232 | 1.386 | 0.122 | 0.439 |
|  | Skate spp.-3Ps | 0.114 | 1.772 | 0.037 | 0.350 |
|  | White Hake-3Ps | 0.202 | 1.901 | 0.057 | 0.709 |
|  | Witch Flounder-3Ps | 0.151 | 1.713 | 0.053 | 0.433 |
| B0 | American Plaice-3Ps | 37.855 | 1.304 | 22.486 | 63.728 |
|  | Atlantic Cod-3Ps | 217.244 | 1.520 | 95.601 | 493.668 |
|  | Haddock-3Ps | 12.075 | 1.583 | 4.910 | 29.693 |
|  | Redfish spp.-3Ps | 160.871 | 1.505 | 72.210 | 358.393 |
|  | Skate spp.-3Ps | 39.540 | 1.819 | 12.239 | 127.742 |
|  | White Hake-3Ps | 15.628 | 1.767 | 5.121 | 47.697 |
|  | Witch Flounder-3Ps | 22.418 | 1.467 | 10.575 | 47.527 |
| sd\_B | American Plaice-3Ps | 0.007 | 4.525 | 0.000 | 0.135 |
|  | Atlantic Cod-3Ps | 0.009 | 4.375 | 0.001 | 0.165 |
|  | Haddock-3Ps | 0.012 | 4.035 | 0.001 | 0.179 |
|  | Redfish spp.-3Ps | 0.072 | 1.822 | 0.022 | 0.232 |
|  | Skate spp.-3Ps | 0.088 | 1.447 | 0.043 | 0.182 |
|  | White Hake-3Ps | 0.187 | 1.486 | 0.086 | 0.408 |
|  | Witch Flounder-3Ps | 0.038 | 2.120 | 0.009 | 0.164 |
| rho | Atlantic Cod-3Ps - American Plaice-3Ps | 0.660 | 0.767 | -0.832 | 0.992 |
|  | Haddock-3Ps - American Plaice-3Ps | 0.497 | 0.795 | -0.919 | 0.991 |
|  | Redfish spp.-3Ps - American Plaice-3Ps | 0.523 | 0.813 | -0.928 | 0.993 |
|  | Skate spp.-3Ps - American Plaice-3Ps | 0.563 | 0.701 | -0.789 | 0.982 |
|  | White Hake-3Ps - American Plaice-3Ps | 0.581 | 0.713 | -0.795 | 0.984 |
|  | Witch Flounder-3Ps - American Plaice-3Ps | 0.633 | 0.701 | -0.743 | 0.985 |
|  | Haddock-3Ps - Atlantic Cod-3Ps | 0.603 | 0.735 | -0.815 | 0.988 |
|  | Redfish spp.-3Ps - Atlantic Cod-3Ps | 0.536 | 0.755 | -0.869 | 0.987 |
|  | Skate spp.-3Ps - Atlantic Cod-3Ps | 0.626 | 0.729 | -0.793 | 0.988 |
|  | White Hake-3Ps - Atlantic Cod-3Ps | 0.730 | 0.712 | -0.674 | 0.991 |
|  | Witch Flounder-3Ps - Atlantic Cod-3Ps | 0.581 | 0.753 | -0.850 | 0.989 |
|  | Redfish spp.-3Ps - Haddock-3Ps | 0.663 | 0.752 | -0.806 | 0.991 |
|  | Skate spp.-3Ps - Haddock-3Ps | 0.668 | 0.734 | -0.774 | 0.990 |
|  | White Hake-3Ps - Haddock-3Ps | 0.792 | 0.692 | -0.534 | 0.992 |
|  | Witch Flounder-3Ps - Haddock-3Ps | 0.210 | 0.854 | -0.979 | 0.991 |
|  | Skate spp.-3Ps - Redfish spp.-3Ps | 0.552 | 0.752 | -0.861 | 0.988 |
|  | White Hake-3Ps - Redfish spp.-3Ps | 0.524 | 0.781 | -0.900 | 0.990 |
|  | Witch Flounder-3Ps - Redfish spp.-3Ps | 0.593 | 0.755 | -0.847 | 0.989 |
|  | White Hake-3Ps - Skate spp.-3Ps | 0.596 | 0.764 | -0.857 | 0.990 |
|  | Witch Flounder-3Ps - Skate spp.-3Ps | 0.637 | 0.767 | -0.843 | 0.992 |
|  | Witch Flounder-3Ps - White Hake-3Ps | 0.394 | 0.853 | -0.968 | 0.994 |
| phi | All species | 0.227 | 0.761 | 0.029 | 0.740 |
| q | American Plaice-3Ps-Spring-Campelen | 1.435 | 1.379 | 0.764 | 2.696 |
|  | American Plaice-3Ps-Spring-Engel | 1.182 | 1.255 | 0.757 | 1.845 |
|  | American Plaice-3Ps-Spring-Yankee | 0.361 | 1.288 | 0.220 | 0.593 |
|  | Atlantic Cod-3Ps-Spring-Campelen | 0.374 | 1.399 | 0.194 | 0.722 |
|  | Atlantic Cod-3Ps-Spring-Engel | 0.307 | 1.399 | 0.159 | 0.593 |
|  | Atlantic Cod-3Ps-Spring-Yankee | 0.056 | 1.453 | 0.027 | 0.117 |
|  | Haddock-3Ps-Spring-Campelen | 0.277 | 1.537 | 0.119 | 0.643 |
|  | Haddock-3Ps-Spring-Engel | 0.364 | 1.444 | 0.177 | 0.747 |
|  | Haddock-3Ps-Spring-Yankee | 0.049 | 1.464 | 0.023 | 0.103 |
|  | Redfish spp.-3Ps-Spring-Campelen | 0.176 | 1.346 | 0.098 | 0.315 |
|  | Redfish spp.-3Ps-Spring-Engel | 0.224 | 1.580 | 0.091 | 0.549 |
|  | Redfish spp.-3Ps-Spring-Yankee | 0.146 | 1.633 | 0.056 | 0.381 |
|  | Skate spp.-3Ps-Spring-Campelen | 0.454 | 1.645 | 0.171 | 1.204 |
|  | Skate spp.-3Ps-Spring-Engel | 0.206 | 1.652 | 0.077 | 0.552 |
|  | Skate spp.-3Ps-Spring-Yankee | 0.154 | 1.698 | 0.055 | 0.436 |
|  | White Hake-3Ps-Spring-Campelen | 0.656 | 1.637 | 0.250 | 1.723 |
|  | White Hake-3Ps-Spring-Engel | 0.208 | 1.712 | 0.072 | 0.596 |
|  | White Hake-3Ps-Spring-Yankee | 0.151 | 1.627 | 0.058 | 0.393 |
|  | Witch Flounder-3Ps-Spring-Campelen | 0.620 | 1.514 | 0.275 | 1.398 |
|  | Witch Flounder-3Ps-Spring-Engel | 0.266 | 1.576 | 0.109 | 0.647 |
|  | Witch Flounder-3Ps-Spring-Yankee | 0.101 | 1.569 | 0.042 | 0.245 |
| sd\_I | American Plaice-3Ps-Spring-Campelen | 0.203 | 1.159 | 0.152 | 0.271 |
|  | American Plaice-3Ps-Spring-Engel | 0.398 | 1.274 | 0.247 | 0.639 |
|  | American Plaice-3Ps-Spring-Yankee | 0.310 | 1.246 | 0.201 | 0.476 |
|  | Atlantic Cod-3Ps-Spring-Campelen | 0.462 | 1.161 | 0.345 | 0.619 |
|  | Atlantic Cod-3Ps-Spring-Engel | 0.650 | 1.232 | 0.432 | 0.979 |
|  | Atlantic Cod-3Ps-Spring-Yankee | 0.469 | 1.247 | 0.304 | 0.724 |
|  | Haddock-3Ps-Spring-Campelen | 0.797 | 1.156 | 0.600 | 1.059 |
|  | Haddock-3Ps-Spring-Engel | 0.746 | 1.235 | 0.494 | 1.128 |
|  | Haddock-3Ps-Spring-Yankee | 0.560 | 1.204 | 0.390 | 0.806 |
|  | Redfish spp.-3Ps-Spring-Campelen | 0.574 | 1.161 | 0.428 | 0.770 |
|  | Redfish spp.-3Ps-Spring-Engel | 0.373 | 1.251 | 0.241 | 0.579 |
|  | Redfish spp.-3Ps-Spring-Yankee | 0.486 | 1.249 | 0.314 | 0.751 |
|  | Skate spp.-3Ps-Spring-Campelen | 0.194 | 1.186 | 0.139 | 0.272 |
|  | Skate spp.-3Ps-Spring-Engel | 0.244 | 1.254 | 0.156 | 0.380 |
|  | Skate spp.-3Ps-Spring-Yankee | 0.497 | 1.289 | 0.302 | 0.819 |
|  | White Hake-3Ps-Spring-Campelen | 0.218 | 1.258 | 0.139 | 0.342 |
|  | White Hake-3Ps-Spring-Engel | 0.203 | 1.866 | 0.060 | 0.691 |
|  | White Hake-3Ps-Spring-Yankee | 0.634 | 1.302 | 0.378 | 1.063 |
|  | Witch Flounder-3Ps-Spring-Campelen | 0.288 | 1.167 | 0.213 | 0.390 |
|  | Witch Flounder-3Ps-Spring-Engel | 0.267 | 1.257 | 0.171 | 0.418 |
|  | Witch Flounder-3Ps-Spring-Yankee | 0.568 | 1.264 | 0.359 | 0.899 |

### Population trends

#### Row

##### Observed and predicted index

##### Process error

##### Standardized process error

> Standardized process error = process error, in log space, divided by
> the standard deviation of the process.

##### Process error correlation

##### Biomass

##### Total biomass

##### Harvest rate
