## Supplement 1 for "Summing the parts: Improving population estimates using a state-space multispecies production model": Single-species - Grand Bank.html

multispic diagnostics and results


multispic diagnostics and results

### Inputs

#### Row

##### Index

##### Landings

##### Covariates

N/A; no process error or K covariates supplied

### Residuals

#### Row

##### Residuals ~ predicted value

##### ~ year

##### ~ survey

##### ~ species

### Parameters

#### Row

##### Correlation

##### log(K)

##### log(r)

##### log(B0)

##### log(SD_B_)

##### logit(rho)

N/A; correlation of process errors across species was not
estimated.

##### logit(phi)

N/A; correlation of process errors across time was not estimated.

##### log(q)

##### log(SD_I_)

##### beta_pe_

N/A; no process error covariates supplied

##### beta_K_

N/A; no K covariates supplied

##### Estimates (plot)

##### Estimates (table)

| parameter | group | estimate | CV | lower | upper |
| --- | --- | --- | --- | --- | --- |
| K | American Plaice-3LNO | 560.253 | 1.770 | 182.980 | 1715.397 |
|  | Atlantic Cod-3LNO | 2323.056 | 3.391 | 212.151 | 25437.488 |
|  | Greenland Halibut-3LNO | 113.302 | 1.220 | 76.745 | 167.274 |
|  | Haddock-3LNO | 19.861 | 1.114 | 16.061 | 24.560 |
|  | Redfish spp.-3LNO | 511.569 | 1.695 | 181.942 | 1438.385 |
|  | Skate spp.-3LNO | 198.914 | 1.234 | 131.827 | 300.140 |
|  | Yellowtail Flounder-3LNO | 282.905 | 1.248 | 183.234 | 436.792 |
| r | American Plaice-3LNO | 0.051 | 1.799 | 0.016 | 0.160 |
|  | Atlantic Cod-3LNO | 0.121 | 1.558 | 0.051 | 0.289 |
|  | Greenland Halibut-3LNO | 0.927 | 1.274 | 0.577 | 1.490 |
|  | Haddock-3LNO | 0.887 | 1.194 | 0.627 | 1.255 |
|  | Redfish spp.-3LNO | 0.291 | 1.366 | 0.158 | 0.536 |
|  | Skate spp.-3LNO | 0.210 | 1.316 | 0.122 | 0.359 |
|  | Yellowtail Flounder-3LNO | 0.208 | 1.460 | 0.099 | 0.435 |
| B0 | American Plaice-3LNO | 910.863 | 1.500 | 411.272 | 2017.327 |
|  | Atlantic Cod-3LNO | 470.033 | 1.570 | 194.089 | 1138.298 |
|  | Greenland Halibut-3LNO | 41.193 | 1.770 | 13.458 | 126.085 |
|  | Haddock-3LNO | 0.620 | 1.787 | 0.199 | 1.934 |
|  | Redfish spp.-3LNO | 206.248 | 2.025 | 51.711 | 822.609 |
|  | Skate spp.-3LNO | 472.309 | 1.431 | 233.945 | 953.538 |
|  | Yellowtail Flounder-3LNO | 263.280 | 1.475 | 122.884 | 564.077 |
| sd\_B | American Plaice-3LNO | 0.214 | 1.167 | 0.158 | 0.290 |
|  | Atlantic Cod-3LNO | 0.307 | 1.269 | 0.192 | 0.490 |
|  | Greenland Halibut-3LNO | 0.295 | 1.180 | 0.214 | 0.408 |
|  | Haddock-3LNO | 0.003 | 3.579 | 0.000 | 0.035 |
|  | Redfish spp.-3LNO | 0.205 | 1.343 | 0.115 | 0.366 |
|  | Skate spp.-3LNO | 0.164 | 1.165 | 0.122 | 0.221 |
|  | Yellowtail Flounder-3LNO | 0.145 | 1.267 | 0.091 | 0.230 |
| q | American Plaice-3LNO-Fall-Campelen | 1.783 | 1.287 | 1.088 | 2.922 |
|  | American Plaice-3LNO-Fall-Engel | 0.461 | 1.313 | 0.270 | 0.785 |
|  | American Plaice-3LNO-Spring-Campelen | 1.112 | 1.290 | 0.675 | 1.833 |
|  | American Plaice-3LNO-Spring-Engel | 0.284 | 1.322 | 0.164 | 0.491 |
|  | American Plaice-3LNO-Spring-Yankee | 0.280 | 1.411 | 0.143 | 0.550 |
|  | Atlantic Cod-3LNO-Fall-Campelen | 1.034 | 1.283 | 0.634 | 1.684 |
|  | Atlantic Cod-3LNO-Fall-Engel | 0.677 | 1.337 | 0.383 | 1.197 |
|  | Atlantic Cod-3LNO-Spring-Campelen | 0.890 | 1.289 | 0.541 | 1.463 |
|  | Atlantic Cod-3LNO-Spring-Engel | 0.650 | 1.292 | 0.394 | 1.074 |
|  | Atlantic Cod-3LNO-Spring-Yankee | 0.162 | 1.386 | 0.085 | 0.307 |
|  | Greenland Halibut-3LNO-Fall-Campelen | 0.268 | 1.266 | 0.169 | 0.426 |
|  | Greenland Halibut-3LNO-Fall-Engel | 0.094 | 1.250 | 0.060 | 0.145 |
|  | Greenland Halibut-3LNO-Spring-Campelen | 0.195 | 1.281 | 0.120 | 0.317 |
|  | Greenland Halibut-3LNO-Spring-Engel | 0.043 | 1.342 | 0.024 | 0.077 |
|  | Greenland Halibut-3LNO-Spring-Yankee | 0.040 | 1.447 | 0.019 | 0.083 |
|  | Haddock-3LNO-Fall-Campelen | 0.306 | 1.203 | 0.213 | 0.440 |
|  | Haddock-3LNO-Fall-Engel | 0.712 | 1.374 | 0.382 | 1.328 |
|  | Haddock-3LNO-Spring-Campelen | 0.574 | 1.183 | 0.413 | 0.799 |
|  | Haddock-3LNO-Spring-Engel | 1.206 | 1.291 | 0.731 | 1.991 |
|  | Haddock-3LNO-Spring-Yankee | 0.470 | 1.659 | 0.174 | 1.266 |
|  | Redfish spp.-3LNO-Fall-Campelen | 0.710 | 1.694 | 0.253 | 1.996 |
|  | Redfish spp.-3LNO-Fall-Engel | 0.216 | 1.581 | 0.088 | 0.529 |
|  | Redfish spp.-3LNO-Spring-Campelen | 0.642 | 1.700 | 0.227 | 1.817 |
|  | Redfish spp.-3LNO-Spring-Engel | 0.174 | 1.480 | 0.081 | 0.376 |
|  | Redfish spp.-3LNO-Spring-Yankee | 0.091 | 1.890 | 0.026 | 0.318 |
|  | Skate spp.-3LNO-Fall-Campelen | 1.353 | 1.220 | 0.916 | 1.999 |
|  | Skate spp.-3LNO-Fall-Engel | 0.542 | 1.251 | 0.349 | 0.840 |
|  | Skate spp.-3LNO-Spring-Campelen | 0.912 | 1.223 | 0.615 | 1.353 |
|  | Skate spp.-3LNO-Spring-Engel | 0.309 | 1.251 | 0.199 | 0.479 |
|  | Skate spp.-3LNO-Spring-Yankee | 0.268 | 1.357 | 0.147 | 0.488 |
|  | Yellowtail Flounder-3LNO-Fall-Campelen | 1.225 | 1.253 | 0.788 | 1.906 |
|  | Yellowtail Flounder-3LNO-Fall-Engel | 0.466 | 1.293 | 0.281 | 0.771 |
|  | Yellowtail Flounder-3LNO-Spring-Campelen | 1.193 | 1.256 | 0.763 | 1.865 |
|  | Yellowtail Flounder-3LNO-Spring-Engel | 0.491 | 1.281 | 0.302 | 0.798 |
|  | Yellowtail Flounder-3LNO-Spring-Yankee | 0.390 | 1.386 | 0.205 | 0.738 |
| sd\_I | American Plaice-3LNO-Fall-Campelen | 0.084 | 1.502 | 0.038 | 0.187 |
|  | American Plaice-3LNO-Fall-Engel | 0.090 | 1.247 | 0.058 | 0.139 |
|  | American Plaice-3LNO-Spring-Campelen | 0.281 | 1.169 | 0.207 | 0.382 |
|  | American Plaice-3LNO-Spring-Engel | 0.182 | 1.571 | 0.075 | 0.442 |
|  | American Plaice-3LNO-Spring-Yankee | 0.073 | 1.565 | 0.030 | 0.176 |
|  | Atlantic Cod-3LNO-Fall-Campelen | 0.282 | 1.235 | 0.186 | 0.426 |
|  | Atlantic Cod-3LNO-Fall-Engel | 0.221 | 1.402 | 0.114 | 0.428 |
|  | Atlantic Cod-3LNO-Spring-Campelen | 0.432 | 1.194 | 0.305 | 0.612 |
|  | Atlantic Cod-3LNO-Spring-Engel | 0.399 | 1.296 | 0.240 | 0.664 |
|  | Atlantic Cod-3LNO-Spring-Yankee | 0.133 | 1.673 | 0.049 | 0.365 |
|  | Greenland Halibut-3LNO-Fall-Campelen | 0.091 | 1.842 | 0.027 | 0.301 |
|  | Greenland Halibut-3LNO-Fall-Engel | 0.132 | 3.553 | 0.011 | 1.579 |
|  | Greenland Halibut-3LNO-Spring-Campelen | 0.372 | 1.175 | 0.271 | 0.511 |
|  | Greenland Halibut-3LNO-Spring-Engel | 0.610 | 1.304 | 0.363 | 1.027 |
|  | Greenland Halibut-3LNO-Spring-Yankee | 0.371 | 1.507 | 0.166 | 0.830 |
|  | Haddock-3LNO-Fall-Campelen | 0.881 | 1.152 | 0.668 | 1.163 |
|  | Haddock-3LNO-Fall-Engel | 0.750 | 1.373 | 0.403 | 1.396 |
|  | Haddock-3LNO-Spring-Campelen | 0.745 | 1.162 | 0.555 | 1.001 |
|  | Haddock-3LNO-Spring-Engel | 0.828 | 1.227 | 0.555 | 1.236 |
|  | Haddock-3LNO-Spring-Yankee | 0.955 | 1.304 | 0.567 | 1.607 |
|  | Redfish spp.-3LNO-Fall-Campelen | 0.385 | 1.196 | 0.271 | 0.546 |
|  | Redfish spp.-3LNO-Fall-Engel | 0.667 | 1.374 | 0.358 | 1.244 |
|  | Redfish spp.-3LNO-Spring-Campelen | 0.489 | 1.182 | 0.352 | 0.679 |
|  | Redfish spp.-3LNO-Spring-Engel | 0.970 | 1.241 | 0.635 | 1.482 |
|  | Redfish spp.-3LNO-Spring-Yankee | 1.095 | 1.363 | 0.596 | 2.010 |
|  | Skate spp.-3LNO-Fall-Campelen | 0.146 | 1.251 | 0.094 | 0.227 |
|  | Skate spp.-3LNO-Fall-Engel | 0.080 | 1.448 | 0.039 | 0.165 |
|  | Skate spp.-3LNO-Spring-Campelen | 0.170 | 1.201 | 0.119 | 0.244 |
|  | Skate spp.-3LNO-Spring-Engel | 0.120 | 1.617 | 0.047 | 0.308 |
|  | Skate spp.-3LNO-Spring-Yankee | 0.096 | 1.661 | 0.035 | 0.259 |
|  | Yellowtail Flounder-3LNO-Fall-Campelen | 0.165 | 1.215 | 0.112 | 0.242 |
|  | Yellowtail Flounder-3LNO-Fall-Engel | 0.225 | 1.249 | 0.145 | 0.348 |
|  | Yellowtail Flounder-3LNO-Spring-Campelen | 0.237 | 1.170 | 0.174 | 0.323 |
|  | Yellowtail Flounder-3LNO-Spring-Engel | 0.176 | 1.310 | 0.104 | 0.298 |
|  | Yellowtail Flounder-3LNO-Spring-Yankee | 0.153 | 1.412 | 0.078 | 0.302 |

### Population trends

#### Row

##### Observed and predicted index

##### Process error

##### Standardized process error

> Standardized process error = process error, in log space, divided by
> the standard deviation of the process.

##### Process error correlation

N/A; correlation of process errors across species was not
estimated.

##### Biomass

##### Total biomass

##### Harvest rate
