## Supplement 1 for "Summing the parts: Improving population estimates using a state-space multispecies production model": Single-species - Northeast NL Shelf.html

multispic diagnostics and results


multispic diagnostics and results

### Inputs

#### Row

##### Index

##### Landings

##### Covariates

N/A; no process error or K covariates supplied

### Residuals

#### Row

##### Residuals ~ predicted value

##### ~ year

##### ~ survey

##### ~ species

### Parameters

#### Row

##### Correlation

##### log(K)

##### log(r)

##### log(B0)

##### log(SD_B_)

##### logit(rho)

N/A; correlation of process errors across species was not
estimated.

##### logit(phi)

N/A; correlation of process errors across time was not estimated.

##### log(q)

##### log(SD_I_)

##### beta_pe_

N/A; no process error covariates supplied

##### beta_K_

N/A; no K covariates supplied

##### Estimates (plot)

##### Estimates (table)

| parameter | group | estimate | CV | lower | upper |
| --- | --- | --- | --- | --- | --- |
| K | American Plaice-2J3K | 75.456 | 1.784 | 24.251 | 234.779 |
|  | Atlantic Cod-2J3K | 1096.847 | 1.823 | 338.109 | 3558.239 |
|  | Greenland Halibut-2J3K | 303.311 | 1.701 | 107.019 | 859.641 |
|  | Redfish spp.-2J3K | 352.964 | 1.909 | 99.360 | 1253.859 |
|  | Skate spp.-2J3K | 17.458 | 1.454 | 8.385 | 36.345 |
|  | Witch Flounder-2J3K | 42.473 | 1.529 | 18.469 | 97.675 |
|  | Wolffish spp.-2J3K | 27.083 | 1.768 | 8.863 | 82.754 |
| r | American Plaice-2J3K | 0.076 | 1.827 | 0.023 | 0.247 |
|  | Atlantic Cod-2J3K | 0.239 | 1.289 | 0.145 | 0.393 |
|  | Greenland Halibut-2J3K | 0.116 | 1.836 | 0.035 | 0.380 |
|  | Redfish spp.-2J3K | 0.172 | 2.056 | 0.042 | 0.705 |
|  | Skate spp.-2J3K | 0.111 | 1.712 | 0.039 | 0.319 |
|  | Witch Flounder-2J3K | 0.150 | 1.540 | 0.064 | 0.350 |
|  | Wolffish spp.-2J3K | 0.095 | 1.972 | 0.025 | 0.359 |
| B0 | American Plaice-2J3K | 226.749 | 1.617 | 88.393 | 581.661 |
|  | Atlantic Cod-2J3K | 522.454 | 1.572 | 215.207 | 1268.352 |
|  | Greenland Halibut-2J3K | 621.953 | 1.866 | 183.133 | 2112.264 |
|  | Redfish spp.-2J3K | 992.329 | 2.512 | 163.107 | 6037.251 |
|  | Skate spp.-2J3K | 33.335 | 1.458 | 15.914 | 69.828 |
|  | Witch Flounder-2J3K | 36.726 | 1.405 | 18.868 | 71.485 |
|  | Wolffish spp.-2J3K | 71.791 | 1.672 | 26.209 | 196.651 |
| sd\_B | American Plaice-2J3K | 0.333 | 1.127 | 0.263 | 0.421 |
|  | Atlantic Cod-2J3K | 0.206 | 1.524 | 0.090 | 0.469 |
|  | Greenland Halibut-2J3K | 0.246 | 1.151 | 0.187 | 0.324 |
|  | Redfish spp.-2J3K | 0.554 | 1.228 | 0.371 | 0.829 |
|  | Skate spp.-2J3K | 0.197 | 1.158 | 0.148 | 0.263 |
|  | Witch Flounder-2J3K | 0.306 | 1.165 | 0.227 | 0.413 |
|  | Wolffish spp.-2J3K | 0.385 | 1.145 | 0.295 | 0.503 |
| q | American Plaice-2J3K-Fall-Campelen | 1.072 | 1.392 | 0.560 | 2.050 |
|  | American Plaice-2J3K-Fall-Engel | 0.445 | 1.416 | 0.225 | 0.881 |
|  | Atlantic Cod-2J3K-Fall-Campelen | 0.816 | 1.415 | 0.413 | 1.612 |
|  | Atlantic Cod-2J3K-Fall-Engel | 0.813 | 1.253 | 0.522 | 1.264 |
|  | Greenland Halibut-2J3K-Fall-Campelen | 0.846 | 1.798 | 0.268 | 2.671 |
|  | Greenland Halibut-2J3K-Fall-Engel | 0.267 | 1.779 | 0.086 | 0.824 |
|  | Redfish spp.-2J3K-Fall-Campelen | 0.692 | 1.890 | 0.199 | 2.410 |
|  | Redfish spp.-2J3K-Fall-Engel | 0.294 | 2.005 | 0.075 | 1.150 |
|  | Skate spp.-2J3K-Fall-Campelen | 1.086 | 1.358 | 0.596 | 1.977 |
|  | Skate spp.-2J3K-Fall-Engel | 0.418 | 1.366 | 0.227 | 0.771 |
|  | Witch Flounder-2J3K-Fall-Campelen | 1.034 | 1.398 | 0.536 | 1.994 |
|  | Witch Flounder-2J3K-Fall-Engel | 0.540 | 1.164 | 0.401 | 0.727 |
|  | Wolffish spp.-2J3K-Fall-Campelen | 0.877 | 1.398 | 0.455 | 1.690 |
|  | Wolffish spp.-2J3K-Fall-Engel | 0.603 | 1.422 | 0.302 | 1.202 |
| sd\_I | American Plaice-2J3K-Fall-Campelen | 0.062 | 1.760 | 0.020 | 0.188 |
|  | American Plaice-2J3K-Fall-Engel | 0.099 | 1.632 | 0.038 | 0.258 |
|  | Atlantic Cod-2J3K-Fall-Campelen | 0.174 | 1.538 | 0.075 | 0.406 |
|  | Atlantic Cod-2J3K-Fall-Engel | 0.688 | 1.214 | 0.471 | 1.006 |
|  | Greenland Halibut-2J3K-Fall-Campelen | 0.047 | 2.006 | 0.012 | 0.184 |
|  | Greenland Halibut-2J3K-Fall-Engel | 0.119 | 1.600 | 0.047 | 0.298 |
|  | Redfish spp.-2J3K-Fall-Campelen | 0.090 | 4.048 | 0.006 | 1.400 |
|  | Redfish spp.-2J3K-Fall-Engel | 0.552 | 1.747 | 0.185 | 1.648 |
|  | Skate spp.-2J3K-Fall-Campelen | 0.080 | 1.371 | 0.043 | 0.149 |
|  | Skate spp.-2J3K-Fall-Engel | 0.094 | 1.439 | 0.046 | 0.192 |
|  | Witch Flounder-2J3K-Fall-Campelen | 0.143 | 1.704 | 0.050 | 0.406 |
|  | Witch Flounder-2J3K-Fall-Engel | 0.045 | 2.488 | 0.008 | 0.268 |
|  | Wolffish spp.-2J3K-Fall-Campelen | 0.132 | 1.666 | 0.048 | 0.358 |
|  | Wolffish spp.-2J3K-Fall-Engel | 0.070 | 2.072 | 0.017 | 0.292 |

### Population trends

#### Row

##### Observed and predicted index

##### Process error

##### Standardized process error

> Standardized process error = process error, in log space, divided by
> the standard deviation of the process.

##### Process error correlation

N/A; correlation of process errors across species was not
estimated.

##### Biomass

##### Total biomass

##### Harvest rate
