## Supplement 1 for "Summing the parts: Improving population estimates using a state-space multispecies production model": Single-species - Southern NL.html

multispic diagnostics and results


multispic diagnostics and results

### Inputs

#### Row

##### Index

##### Landings

##### Covariates

N/A; no process error or K covariates supplied

### Residuals

#### Row

##### Residuals ~ predicted value

##### ~ year

##### ~ survey

##### ~ species

### Parameters

#### Row

##### Correlation

##### log(K)

##### log(r)

##### log(B0)

##### log(SD_B_)

##### logit(rho)

N/A; correlation of process errors across species was not
estimated.

##### logit(phi)

N/A; correlation of process errors across time was not estimated.

##### log(q)

##### log(SD_I_)

##### beta_pe_

N/A; no process error covariates supplied

##### beta_K_

N/A; no K covariates supplied

##### Estimates (plot)

##### Estimates (table)

| parameter | group | estimate | CV | lower | upper |
| --- | --- | --- | --- | --- | --- |
| K | American Plaice-3Ps | 48.955 | 1.256 | 31.299 | 76.571 |
|  | Atlantic Cod-3Ps | 208.476 | 1.359 | 114.316 | 380.193 |
|  | Haddock-3Ps | 17.044 | 1.128 | 13.451 | 21.598 |
|  | Redfish spp.-3Ps | 173.638 | 1.683 | 62.565 | 481.900 |
|  | Skate spp.-3Ps | 32.962 | 1.580 | 13.446 | 80.805 |
|  | White Hake-3Ps | 23.541 | 1.539 | 10.115 | 54.790 |
|  | Witch Flounder-3Ps | 18.150 | 1.554 | 7.647 | 43.083 |
| r | American Plaice-3Ps | 0.258 | 1.320 | 0.150 | 0.444 |
|  | Atlantic Cod-3Ps | 0.920 | 1.356 | 0.507 | 1.671 |
|  | Haddock-3Ps | 0.569 | 1.337 | 0.322 | 1.004 |
|  | Redfish spp.-3Ps | 0.260 | 1.952 | 0.070 | 0.965 |
|  | Skate spp.-3Ps | 0.411 | 1.635 | 0.157 | 1.078 |
|  | White Hake-3Ps | 0.201 | 1.698 | 0.071 | 0.567 |
|  | Witch Flounder-3Ps | 0.676 | 1.391 | 0.354 | 1.292 |
| B0 | American Plaice-3Ps | 55.928 | 1.387 | 29.460 | 106.177 |
|  | Atlantic Cod-3Ps | 145.766 | 1.900 | 41.435 | 512.796 |
|  | Haddock-3Ps | 36.111 | 1.397 | 18.748 | 69.551 |
|  | Redfish spp.-3Ps | 114.033 | 1.750 | 38.074 | 341.530 |
|  | Skate spp.-3Ps | 41.258 | 1.920 | 11.491 | 148.134 |
|  | White Hake-3Ps | 11.732 | 1.886 | 3.384 | 40.675 |
|  | Witch Flounder-3Ps | 15.915 | 1.984 | 4.154 | 60.970 |
| sd\_B | American Plaice-3Ps | 0.011 | 3.996 | 0.001 | 0.165 |
|  | Atlantic Cod-3Ps | 0.341 | 1.362 | 0.186 | 0.625 |
|  | Haddock-3Ps | 0.005 | 3.491 | 0.000 | 0.061 |
|  | Redfish spp.-3Ps | 0.025 | 3.976 | 0.002 | 0.378 |
|  | Skate spp.-3Ps | 0.182 | 1.377 | 0.097 | 0.341 |
|  | White Hake-3Ps | 0.297 | 1.251 | 0.192 | 0.461 |
|  | Witch Flounder-3Ps | 0.205 | 1.441 | 0.100 | 0.419 |
| q | American Plaice-3Ps-Spring-Campelen | 0.462 | 1.247 | 0.299 | 0.712 |
|  | American Plaice-3Ps-Spring-Engel | 0.679 | 1.488 | 0.312 | 1.481 |
|  | American Plaice-3Ps-Spring-Yankee | 0.320 | 1.433 | 0.158 | 0.648 |
|  | Atlantic Cod-3Ps-Spring-Campelen | 0.256 | 1.491 | 0.117 | 0.561 |
|  | Atlantic Cod-3Ps-Spring-Engel | 0.340 | 1.625 | 0.131 | 0.880 |
|  | Atlantic Cod-3Ps-Spring-Yankee | 0.104 | 1.547 | 0.044 | 0.244 |
|  | Haddock-3Ps-Spring-Campelen | 0.143 | 1.202 | 0.100 | 0.206 |
|  | Haddock-3Ps-Spring-Engel | 0.698 | 1.444 | 0.340 | 1.434 |
|  | Haddock-3Ps-Spring-Yankee | 0.046 | 1.236 | 0.030 | 0.069 |
|  | Redfish spp.-3Ps-Spring-Campelen | 0.431 | 1.650 | 0.162 | 1.150 |
|  | Redfish spp.-3Ps-Spring-Engel | 0.357 | 1.519 | 0.157 | 0.810 |
|  | Redfish spp.-3Ps-Spring-Yankee | 0.240 | 1.831 | 0.073 | 0.786 |
|  | Skate spp.-3Ps-Spring-Campelen | 0.828 | 1.639 | 0.314 | 2.181 |
|  | Skate spp.-3Ps-Spring-Engel | 0.379 | 1.623 | 0.147 | 0.980 |
|  | Skate spp.-3Ps-Spring-Yankee | 0.206 | 1.678 | 0.075 | 0.569 |
|  | White Hake-3Ps-Spring-Campelen | 0.494 | 1.765 | 0.162 | 1.504 |
|  | White Hake-3Ps-Spring-Engel | 0.228 | 1.641 | 0.086 | 0.602 |
|  | White Hake-3Ps-Spring-Yankee | 0.173 | 1.733 | 0.059 | 0.508 |
|  | Witch Flounder-3Ps-Spring-Campelen | 0.554 | 1.578 | 0.227 | 1.355 |
|  | Witch Flounder-3Ps-Spring-Engel | 0.252 | 1.626 | 0.097 | 0.654 |
|  | Witch Flounder-3Ps-Spring-Yankee | 0.119 | 1.751 | 0.040 | 0.357 |
| sd\_I | American Plaice-3Ps-Spring-Campelen | 0.205 | 1.153 | 0.155 | 0.271 |
|  | American Plaice-3Ps-Spring-Engel | 0.763 | 1.215 | 0.521 | 1.118 |
|  | American Plaice-3Ps-Spring-Yankee | 0.344 | 1.264 | 0.217 | 0.544 |
|  | Atlantic Cod-3Ps-Spring-Campelen | 0.424 | 1.280 | 0.261 | 0.687 |
|  | Atlantic Cod-3Ps-Spring-Engel | 0.500 | 1.413 | 0.254 | 0.985 |
|  | Atlantic Cod-3Ps-Spring-Yankee | 0.299 | 1.436 | 0.147 | 0.608 |
|  | Haddock-3Ps-Spring-Campelen | 0.762 | 1.154 | 0.576 | 1.009 |
|  | Haddock-3Ps-Spring-Engel | 0.831 | 1.231 | 0.553 | 1.249 |
|  | Haddock-3Ps-Spring-Yankee | 0.466 | 1.201 | 0.326 | 0.667 |
|  | Redfish spp.-3Ps-Spring-Campelen | 0.597 | 1.159 | 0.447 | 0.797 |
|  | Redfish spp.-3Ps-Spring-Engel | 0.406 | 1.241 | 0.266 | 0.620 |
|  | Redfish spp.-3Ps-Spring-Yankee | 0.508 | 1.252 | 0.327 | 0.789 |
|  | Skate spp.-3Ps-Spring-Campelen | 0.140 | 1.436 | 0.069 | 0.284 |
|  | Skate spp.-3Ps-Spring-Engel | 0.203 | 1.451 | 0.098 | 0.421 |
|  | Skate spp.-3Ps-Spring-Yankee | 0.414 | 1.283 | 0.254 | 0.674 |
|  | White Hake-3Ps-Spring-Campelen | 0.154 | 1.688 | 0.055 | 0.429 |
|  | White Hake-3Ps-Spring-Engel | 0.121 | 3.009 | 0.014 | 1.051 |
|  | White Hake-3Ps-Spring-Yankee | 0.582 | 1.325 | 0.335 | 1.010 |
|  | Witch Flounder-3Ps-Spring-Campelen | 0.202 | 1.489 | 0.092 | 0.440 |
|  | Witch Flounder-3Ps-Spring-Engel | 0.208 | 1.781 | 0.067 | 0.644 |
|  | Witch Flounder-3Ps-Spring-Yankee | 0.603 | 1.298 | 0.362 | 1.005 |

### Population trends

#### Row

##### Observed and predicted index

##### Process error

##### Standardized process error

> Standardized process error = process error, in log space, divided by
> the standard deviation of the process.

##### Process error correlation

N/A; correlation of process errors across species was not
estimated.

##### Biomass

##### Total biomass

##### Harvest rate
